# Alternative Anastrozole Target CYP4V2 Impairs Neurons in Preclinical Models of Inherited Tauopathy

**DOI:** 10.1101/2025.11.29.689498

**Authors:** Chiara Pedicone, Sarah A. Weitzman, Joshua Orrick, Derian A. Pugh, Alison M. Goate, Ross L. Cagan, Kathryn R. Bowles, Masahiro Sonoshita

**Author notes:** Corresponding authors: Kathryn R Bowles, Masahiro Sonoshita.

## Abstract

Tauopathies, including Alzheimer disease, are neurodegenerative diseases characterized by the pathological aggregation of the microtubule-associated protein Tau (MAPT). This study explores the neuroprotective properties of anastrozole, a clinical Aromatase inhibitor, in both a *Drosophila* tauopathy model expressing human *MAPT*^R406W^ and isogenic pairs of human induced pluripotent stem cell-derived cortical neurons expressing *MAPT* mutations. The Aromatase inhibitor anastrozole, initially identified in a Drosophila *MAPT^R406W^* drug screen, reduced oxidative stress and Tau phosphorylation in cultured cortical neurons. Its protective effects were not mediated by Aromatase activity or cholesterol metabolism: screening in *Drosophila* identified CYP4V2 as a potential off-target. Knockdown of CYP4V2 in *MAPT^R406W^* neurons recapitulated protection against oxidative stress-induced cell death. These findings position anastrozole and its offtarget CYP4V2 as novel therapeutic avenues for primary tauopathies and potentially Alzheimer’s disease and related dementias by reducing Tau phosphorylation and susceptibility to oxidative stress.

## Background

Tauopathies, such as Alzheimer’s disease (AD), progressive supranuclear palsy (PSP), and frontotemporal dementia (FTD), are characterized by the abnormal accumulation of hyperphosphorylated Tau protein within the brain, leading to neuronal loss and cognitive decline ^1^. The microtubule-associated protein Tau, encoded by the *MAPT* gene, plays a key role in maintaining the structural integrity of neurons and promoting intracellular transport processes essential for neuronal function ^1^. Pathogenic mutations such as *MAPT^R406W^* have been shown to promote tauopathies through aggregation of Tau protein, leading to neuronal degeneration ^2^. Despite significant advances in understanding tauopathies, effective therapeutic options remain limited, demonstrating a continuing unmet clinical need.

Tau-targeted therapies, including monoclonal antibodies and small-molecule inhibitors, have shown limited clinical success ^3^. This underscores the need for novel strategies that move beyond Tau clearance alone and address the broader cellular dysfunction associated with tauopathies. Previous work has reported that neurons carrying *MAPT* mutations are more sensitive to oxidative and endoplasmic reticulum stress, suggesting that targeting these cellular responses could mitigate tauopathy-associated neurodegeneration ^4,5^. Given the heightened vulnerability of *MAPT* mutant neurons to cellular stress, therapeutic approaches that restore cellular homeostasis, enhance protein quality control, and modulate stress response pathways could provide greater neuroprotection. Additionally, targeting upstream regulators of Tau pathology may offer more durable interventions.

One useful tool for exploring cell biological processes and therapeutics in the context of the whole body is the fruit fly *Drosophila*, which possesses genes, signaling pathways, and tissue functions similar to those of mammals. In this study, we combine use of a transgenic *Drosophila* model with cultured human induced pluripotent stem cell (iPSC)-derived neurons to identify and validate anastrozole as an effective suppressor of human *MAPT^R406W^*-mediated cellular stress susceptibility. Anastrozole was designed as a non-steroidal Aromatase inhibitor that is commonly used in the treatment of hormone-dependent breast cancer. We demonstrate that anastrozole-mediated protection occurs through an off-target interaction with CYP4V2, in which reduced CYP4V2 expression is sufficient to protect *MAPT^R406W^*neurons from reactive oxygen species (ROS)-mediated cell death. Together, our data support CYP4V2 as a novel candidate therapeutic target for *MAPT*-mutant tauopathies.

## Results

### Drosophila screening identifies anastrozole as a drug candidate for tauopathy

Previous studies have generated *Drosophila* transgenic models using the pan-neuronal driver *elav-gal4* to direct transgenes encoding either wild-type human Tau (*hTau^WT^*) or the R406W mutant isoform associated with familial frontotemporal dementia (*hTau^R406W^*) ^2^. Expression of either construct resulted in neurodegeneration and reduced lifespan in adult flies; *hTau^R406W^* flies exhibited more severe phenotypes. To generate a tauopathy model amenable to chemical screening, we used the driver *GMR-g al4*to express *hTau^WT^* and *hTau^R406W^* specifically in the eye, a useful model for neurodegenerative diseases including tauopathy ^6^. This resulted in a rough eye phenotype indicative of excess neuronal cell death (Fig. 1A). Notably, *GMR>hTau^R406W^* flies also exhibited a reduced eclosion rate (survival), consistent with the shortened longevity previously reported for *hTau^R406W^* versus *hTau^WT^* flies (Fig. 1B) ^2^.

**Figure 1:**
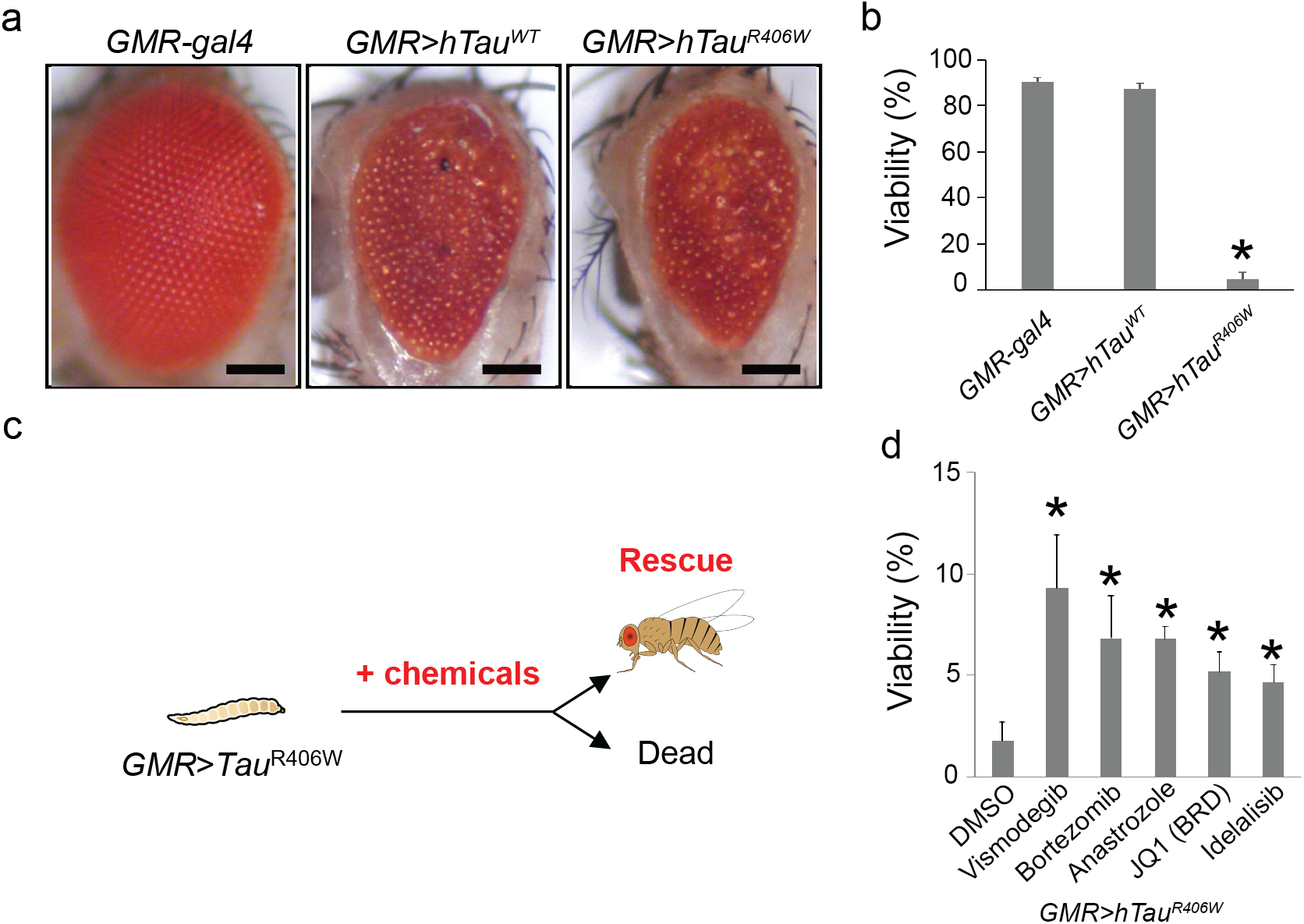
Anastrozole protects from lethality in a MAPT mutant fly model. A) *GMR*>*hTau^WT^* and *GMR*>*hTau^R406W^*flies exhibiting a rough eye phenotype compared to control *GMR-gal4* flies. Scale bars, 100 µm. B) *hTau^R406W^* caused shortened longevity of flies. Fly viability was determined by dividing the number of adults by that of total pupae. Error bars, SD in biological triplicate. \**p* < 0.01 in Dunnet’s test compared with *GMR-gal4*. C) Chemical testing in flies. Vismodegib (Hedgehog pathway inhibitor; final 1 µM in fly food), bortezomib (proteasome inhibitor; 1 µM), anastrozole (Aromatase inhibitor; 100 µM), JQ1 (BET bromodomain inhibitor; 100 µM), and idelalisib (PI3Kδ inhibitor; 200 µM) were each mixed into the food and administered to *GMR*>*hTau^R406W^*flies, respectively. Fly viability was assessed following the same procedure in B. D) Rescue of *GMR*>*hTau^R406W^* lethality by chemicals. Error bars, SD in 8 biological replicates (N > 30 per replicate). \**p* < 0.05 in Dunnet’s test compared with vehicle (DMSO) control.

Next, we leveraged the *GM R>hTau^R406W^*model to screen a library of FDA-approved cancer drugs (Suppl. Table 1) and experimental compounds for chemical modifiers of the rough eye phenotype (Fig. 1C). Mixing compounds in the food for oral delivery, we observed moderate but significant rescue of fly viability with five compounds: vismodegib (Hedgehog pathway SMO inhibitor), bortezomib (26S proteasome inhibitor), anastrozole (Aromatase inhibitor), JQ1 (BET bromodomain inhibitor), and idelalisib (PI3Kδ inhibitor; Fig. 1D).

### Increased stress susceptibility in iPSC-derived MAPT*^R406W^* neurons

To assess these five compounds in a mammalian platform, we used two human isogenic iPSC paired lines derived from patients heterozygous for the *MAPT^R406W^* mutation ^7^ (Suppl. Fig. 1A). Following established protocols, iPSCs were first differentiated into neural progenitor cells (NPCs; Suppl. Fig. 1B), and then further differentiated for 4 weeks into cortical neurons (Suppl. Fig. 1C)^8^.

We established an *in vitro* stress susceptibility assay using a human cortical neuron model to assess our five candidate compounds. Based on reports that *MAPT^A152T^* neurons are susceptible to stressors such as rotenone ^9^, we tested *MAPT^R406W^* neurons for increased susceptibility to rotenone-induced oxidative stress (Suppl. Fig. 1D). Four-week old neurons were incubated for 24 hours with DMSO or rotenone at 500 nM, 1 μM, or 3 μM and subsequently labeled with CellRox Green to quantify ROS. *MAPT^R406W^* neurons showed increased oxidative stress compared to isogenic controls, both at baseline and at all tested rotenone concentrations. ROS levels in *MAPT^R406W^* neurons were higher than controls upon rotenone stimulation (Suppl. Fig. 1D). *MAPT^R406W^* neurons did not show any significant difference in Lactate Dehydrogenase (LDH) release at baseline or after 24 h rotenone treatment (Suppl. Fig. 1E). 500 nM rotenone was used for further drug treatment assays due to its pronounced differential response.

Tau hyperphosphorylation is a key feature of multiple tauopathies ^4,10^. To assess this phenotype in our model, we measured the phosphorylation of three Tau epitopes in *MAPT^R406W^* and control neurons by ELISA—serine 199 (S199), serine 396 (S396) and threonine 181 (T181)—and normalized these measures to total Tau. Similar to previous reports in *MAPT^R406W^* neurons ^11^ we did not observe increased levels of total Tau in 4 week old *MAPT^R406W^* neurons, nor did we observe hyperphosphorylation of Tau (Suppl. Fig. 1F).

### Anastrozole reduced oxidative stress susceptibility in iPSC-derived MAPT*^R406W^* neurons

To test drug efficacy, we first treated neuronal cultures for 24 h with 1 nM to 10 µM of each of the five compound hits and assessed oxidative stress. Minimal toxicity was observed in neurons after 24 h incubation with each compound based on LDH release compared to vehicle (DMSO) control for all concentrations (Suppl. Fig. 2A). The exception was incubation with bortezomib, which led to a significant increase in LDH release at concentrations above 1 µM, suggesting toxicity.

We next assessed the impact of each of the five compounds (1 µM) on rotenone-induced ROS production. Rapamycin was used as a positive control ^4,12^. JQ1 and anastrozole significantly blocked the effect of rotenone on oxidative stress in *MAPT^R406W^* neurons (Suppl. Fig. 2B), while vismodegib and, interestingly, rapamycin proved to be less effective (Suppl. Fig. 2B). In contrast, idelalisib had no effect on oxidative stress, whereas bortezomib significantly increased the level of oxidative stress compared to rotenone alone, consistent with its effect on LDH levels in the absence of a stressor (Suppl. Fig. 2A).

Previous studies have shown that reduced estrogen, particularly 17β-estradiol, is associated with increased risk of tauopathy-related disease, and that 17β-estradiol treatment is neuroprotective and prevents Tau hyperphosphorylation ^13–15^. In contrast, our data indicate that anastrozole, which lowers estrogen levels, protects *MAPT^R406W^* neurons from cellular stress. To investigate its mechanism, we treated *MAPT^R406W^* and isogenic control neurons from two iPSC pairs with anastrozole (10 nM–10 µM) for 24 h before inducing oxidative stress with 500 nM rotenone (Fig. 2A–E). Consistent with our primary screen, anastrozole robustly reduced oxidative stress in *MAPT^R406W^* neurons at all concentrations, such that *MAPT^R406W^* neurons did not differ significantly from isogenic controls for up to 15 h of rotenone exposure (except at 5 µM; Fig. 2F). Anastrozole treatment also significantly decreased S396 phosphorylation in control neurons and T181 phosphorylation in both control and mutant neurons in a dose-dependent manner, without affecting total Tau levels or pS199 in either genotype (Fig. 2G–J). These findings, together with previous studies, raise the question of whether anastrozole’s protective activity in this context is mediated through Aromatase inhibition and consequent reduction of estrogen levels.

**Figure 2:**
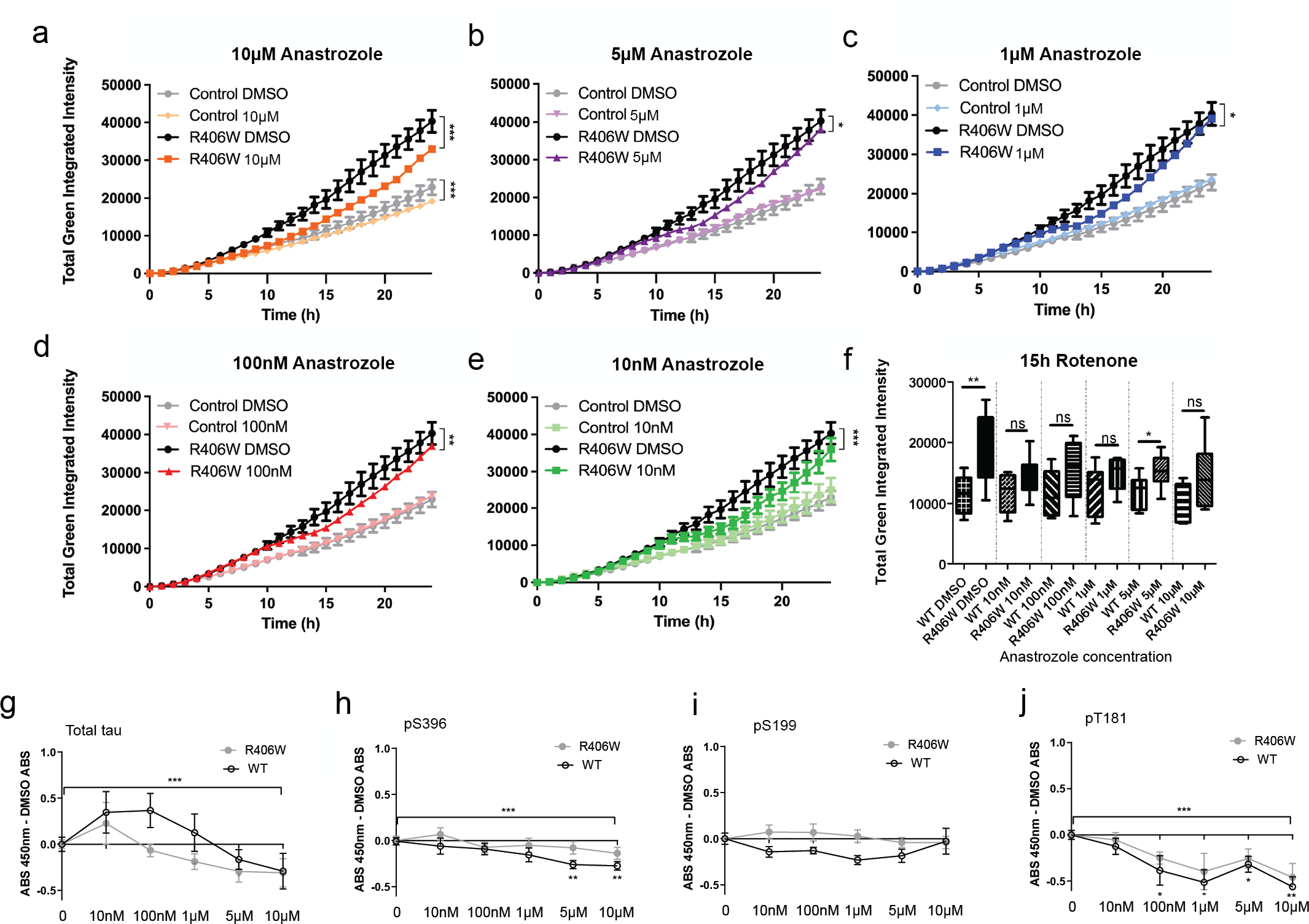
Anastrozole reduces ROS production and Tau phosphorylation in neurons. Quantification of ROS production in neurons over time for different dosages of anastrozole compared to DMSO control, following treatment with rotenone. Linear regression. F) CellRox fluorescence intensity for each anastrozole concentration (A-E) at 15 hours post-rotenone exposure. Student’s *t*-test. A-F N = 2 isogenic pairs, each with three technical replicates per condition consisting of three fields of view (each data point represents a total sample size of N = 18). G-J) Tau phosphorylation in neuronal lysates following 24 hours of anastrozole treatment, relative to DMSO. Data representative of 3 independent experiments on 2 isogenic pairs, with two technical replicates per condition. Two-way ANOVA and Dunnett’s multiple comparisons test. Error bars = SEM. \**p* <0.05, \*\**p* < 0.01, and \*\*\**p* < 0.001. ns, not significant.

### Anastrozole does not protect MAPT*^R406W^* neurons via Aromatase inhibition

To better understand the mechanism of anastrozole’s protective effect, we first asked whether it acts through its canonical target, Aromatase. Anastrozole is a non-steroidal, reversible Aromatase inhibitor used for hormone-dependent breast cancer that blocks CYP19A1-mediated conversion of androgens to estrogens ^16^. We compared anastrozole with an irreversible Aromatase inhibitor (exemestane) and an Estrogen receptor antagonist (fulvestrant) to distinguish effects of reduced estrogen synthesis from downstream receptor signalling. In contrast to anastrozole, exemestane significantly increased oxidative stress in both *MAPT^R406W^* and isogenic control neurons, whereas fulvestrant had no significant effect at any concentration tested (Fig. 3A, B; Suppl. Fig. 3A, B).

**Figure 3:**
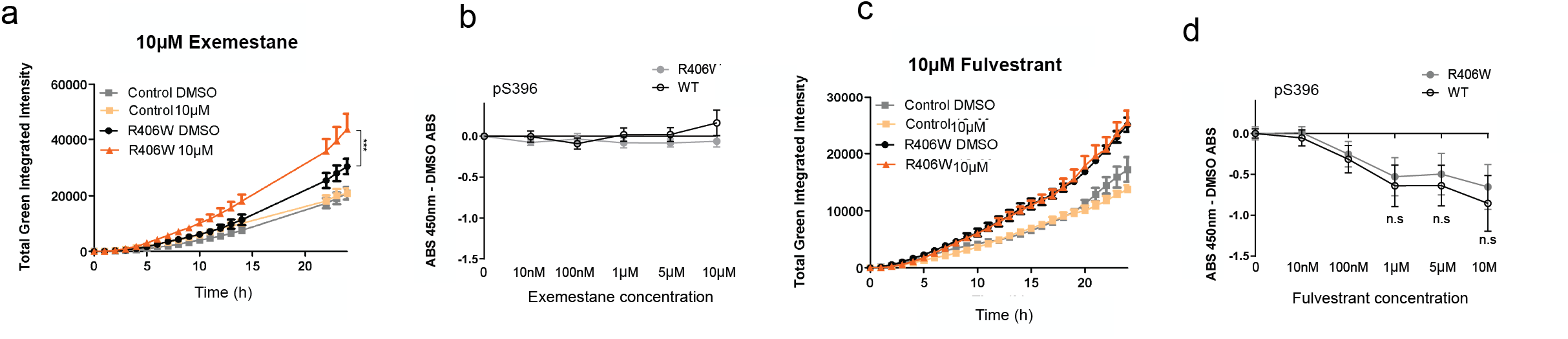
Assessing a role for Aromatase in neuronal protection. A-D) Effect of 10 µM exemestane and 10 µM fulvestrant on ROS production (A-B) and Tau phosphorylation (C-D). Data points represent N = 2 independent isogenic pairs, each consisting of three technical replicates encompassing three fields of view. A-B assessed by linear regression, C-D by two-way ANOVA with Dunnett’s multiple comparisons test. Error bars = SEM. \**p* < 0.05, and \*\*\**p* < 0.001. n.s, not significant.

To assess effects on Tau phosphorylation, we treated *MAPT^R406W^* and control neurons for 24 h with DMSO or 10 nM–10 µM of each compound and quantified phosphorylated Tau by ELISA. Unlike anastrozole, which reduced Tau phosphorylation, exemestane had no significant effect, while fulvestrant induced a mild, dose-dependent decrease in pS396 that did not reach significance (Fig. 3C, 3D, Suppl. Fig. 3C, 3D). The inability of exemestane or fulvestrant to suppress oxidative stress or Tau phosphorylation suggests that anastrozole is unlikely to act through Aromatase inhibition or canonical estrogen signalling. Together with the lack of estrogen-like hormones in *Drosophila*, which showed anastrozole-mediated rescue, these data support an alternative mechanism.

To test this directly, we measured extracellular E2 estradiol and testosterone by ELISA after 24 h of drug treatment. Anastrozole did not alter E2 or testosterone at any concentration at this short time point (Suppl. Fig. 4A, B), indicating that its neuroprotective effect is unlikely to reflect changes in estradiol levels. In contrast, exemestane caused a small increase in E2 and a significant decrease in testosterone, and fulvestrant significantly increased E2 without affecting testosterone (Suppl. Fig. 3A,B). Despite these hormone changes, neither compound rescued *MAPT^R406W^* neurons.

### Cholesterol off-targets of anastrozole do not underlie protection in MAPT*^R406W^* neurons

Previous studies reported that anastrozole can also inhibit the cholesterol oxysterol 27-hydroxycholesterol (27-OHC) via the sterol 27-hydroxylase CYP27A1 ^17,18^. Dysregulated cholesterol metabolism involving CYP46A1 and 24-hydroxycholesterol (24-OHC) has been linked to tauopathy and AD models ^19–21^, and low-dose efavirenz, a CYP46A1 agonist, enhanced cholesterol turnover ^22^. We therefore asked whether modulating cholesterol metabolism would mimic anastrozole’s protection in *MA PT^R406W^* neurons.

We first compared 27-OHC and 24-OHC levels between *MAPT^R406W^* and isogenic control neurons. Both oxysterols were significantly reduced in *MAPT^R406W^* neurons (Fig. 4A), in contrast to reports of elevated 24-OHC and increased CYP46A1 expression in early AD ^23–25^. Moreover, CYP46A1 overexpression is neuroprotective in the THY-Tau22 mouse model ^26^, suggesting that limiting cholesterol accumulation through 24-OHC–dependent turnover can be beneficial in tauopathies. These observations argue against anastrozole acting primarily through cholesterol metabolism in our system. Consistent with this, ELISAs of culture supernatants after 24 h of anastrozole treatment showed no change in 24-OHC or 27-OHC at any concentration (Fig. 4B, C). Neutral triglycerides (Oil Red O) and total cholesterol (Amplex Red) were also similar between *MAPT^R406W^* and control neurons, aside from donor-specific differences (Suppl. Fig. 5A, B).

**Figure 4:**
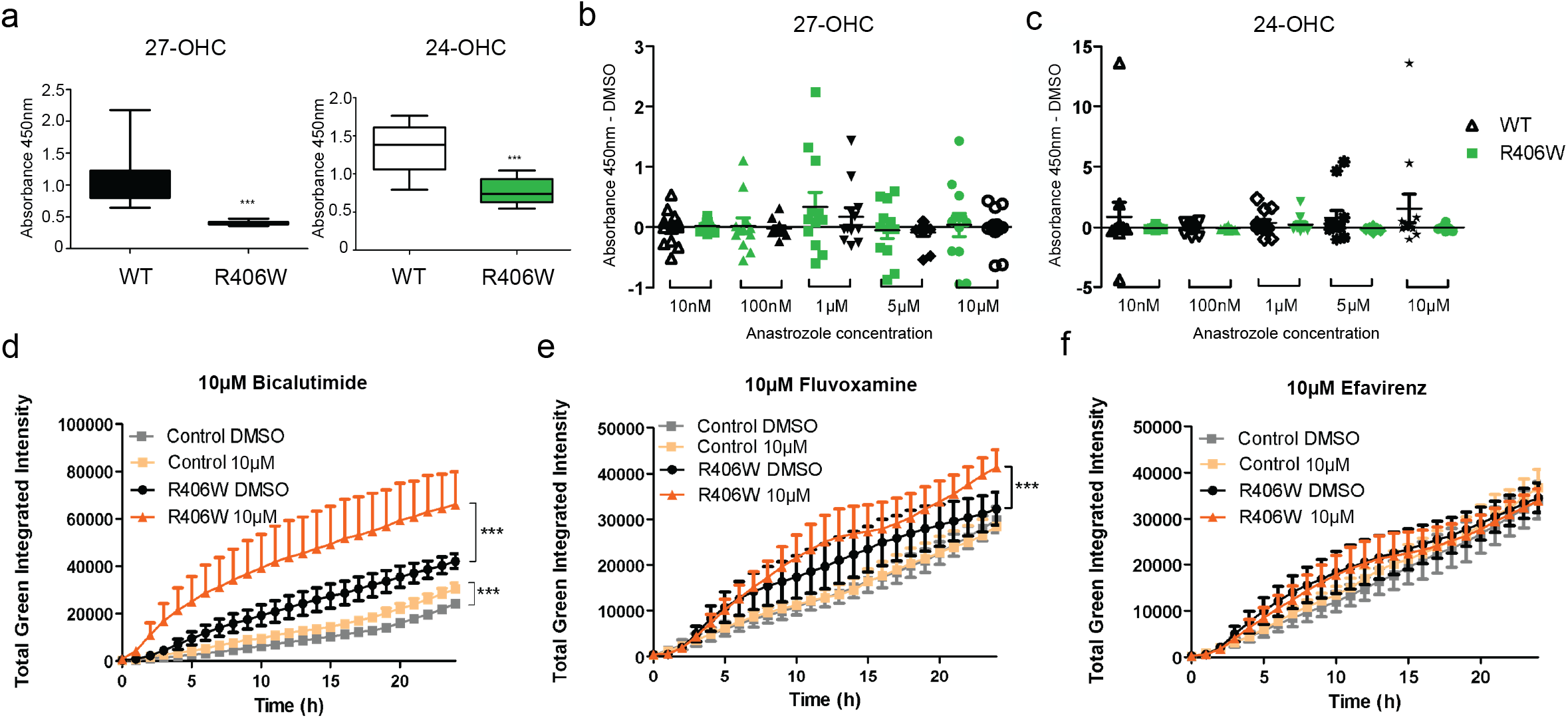
Cholesterol biosynthesis inhibitors did not recapitulate anastrozole protective effect. A) Extracellular levels of 27-OHC and 24-OHC in WT and *MAPT^R406W^*neurons at baseline. Representative data from two independent experiments conducted in two isogenic sets, analyzed using Student’s *t*-test. B-C) Extracellular levels of 27-OHC (B) and 24-OHC (C) following 24h treatment with anastrozole. Representative of two independent experiments on two separate isogenic pairs, encompassing 3 technical replicates per condition. Two-way ANOVA with Dunnett’s multiple comparisons. D-F) Effect of bicalutamide (D), fluvoxamine (E), and efavirenz (F) on ROS production.]Data points represent N = 2 independent isogenic pairs, each consisting of three technical replicates encompassing three fields of view, linear regression. Error bars = SEM. Lower bars removed from D and E for clarity. \*\*\**p* < 0.001.

We next tested other modulators of cholesterol pathways. Bicalutamide (27-OHC inhibitor) modestly lowered levels of Tau^R406W^ at high doses and increased pS199 Tau in controls without reaching significance (Suppl. Fig. 5D). Fluvoxamine (27-OHC activator) did not alter total or phospho-Tau species but, like bicalutamide, increased oxidative stress (Fig. 4D, E; Suppl. Fig. 5C, E). Efavirenz (24-OHC agonist) had no effect on ROS or Tau phosphorylation (Fig. 4F; Suppl. Fig. 5C, F), in contrast to prior AD studies ^21^. ELISAs confirmed the expected reduction in 27-OHC with bicalutamide and revealed unanticipated decreases with efavirenz and fluvoxamine (Suppl. Fig. 6A–C). We conclude that, although cholesterol metabolism is altered in *MAPT^R406W^* neurons, these data do not support it as the mechanism for anastrozole’s protection.

### Anastrozole drives protective transcriptional changes in MAPT*^R406W^* neurons

Anastrozole’s protective effects, independent of Aromatase and CYP27A1, prompted us to perform transcriptomic analysis to screen for alternative mechanisms (Suppl. Table 2,3,4,5). RNA-sequencing of untreated *MAPT^R406W^* neurons identified significant transcriptional dysregulation relative to control neurons (Suppl. Fig. 7A, Suppl. Table 4A). KEGG pathway analysis ranked on q value (Fig. 5A, Suppl. Table 4B) showed upregulation of DNA replication (NES⍰ =⍰ 2.39, q < 10^-5^), mismatch repair (NES⍰ = ⍰1.91, q = 8×10^-3^), and oocyte meiosis (NES ⍰= ⍰1.85, q = 1.2×10^-2^), and downregulation of ribosomal (NES = 2.9, q < 10^-5^) and ABC transporter (NES ⍰=⍰ –1.98, q < 10^-4^) pathways (Table 1).

**Figure 5:**
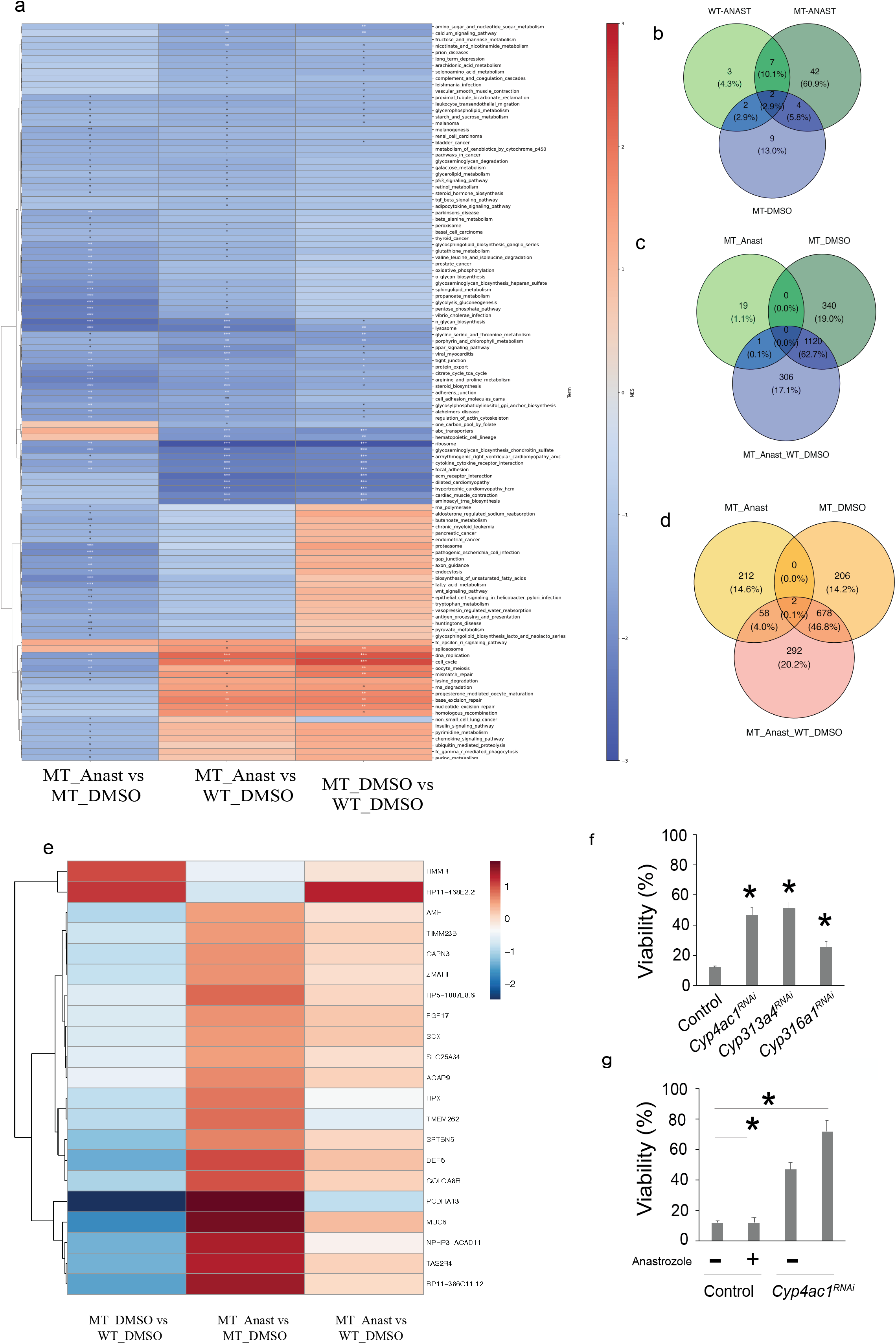
RNA-seq analysis of anastrozole treated WT or MAPT*^R406W^* neurons. A) Comparative pathway analysis showing differential regulation showing anastrozole effect in mutant neurons (MT_Anast vs MT_DMSO), effect of the *MAPT^R406W^* mutation (MT_DMSO vs WT_DMSO), and interaction showing corrective effect of anastrozole in *MAPT^R406W^* compared to WT levels (MT_Anast vs WT_DMSO). Blue bars indicate downregulation and red bars indicate upregulation of pathways, intensity of the scale depends on NES from t-statistics, stars are q values for significance. B) Overlap between significantly downregulated pathways following anastrozole treatment in control (WT_Anast vs. WT_DMSO) and *MAP T^R406W^* (MT_Anast vs. MT_DMSO) neurons, and between *MAPT^R406W^* and control neurons (MT-DMSO vs. WT-DMSO). C-D) Overlap of upregulated (C) and downregulated (D) genes (|log2FC| > 0.5, *p* value < 0.05) between comparisons. (E) Differentially expressed genes downstream of anastrozole treatment in control and *MAPT^R406W^* neurons. F) Targeted RNAi-mediated inhibition of *Cyp4ac1, Cyp313a4*, or *Cyp316a1* rescued lethality of *GM R*>*hTau^R406W^* flies. Error bars, SD in biological triplicate. \**p* < 0.05 in Dunnett’s test compared to control. G) Targeted inhibition of *Cy p4ac1* enhanced the ability of low dose anastrozole treatment (10 µM) to rescue *GMR*>*hTau^R406W^* viability. Error bars, SD in biological triplicate. \**p* < 0.05 in Dunnet’s test.

**Table 1:**
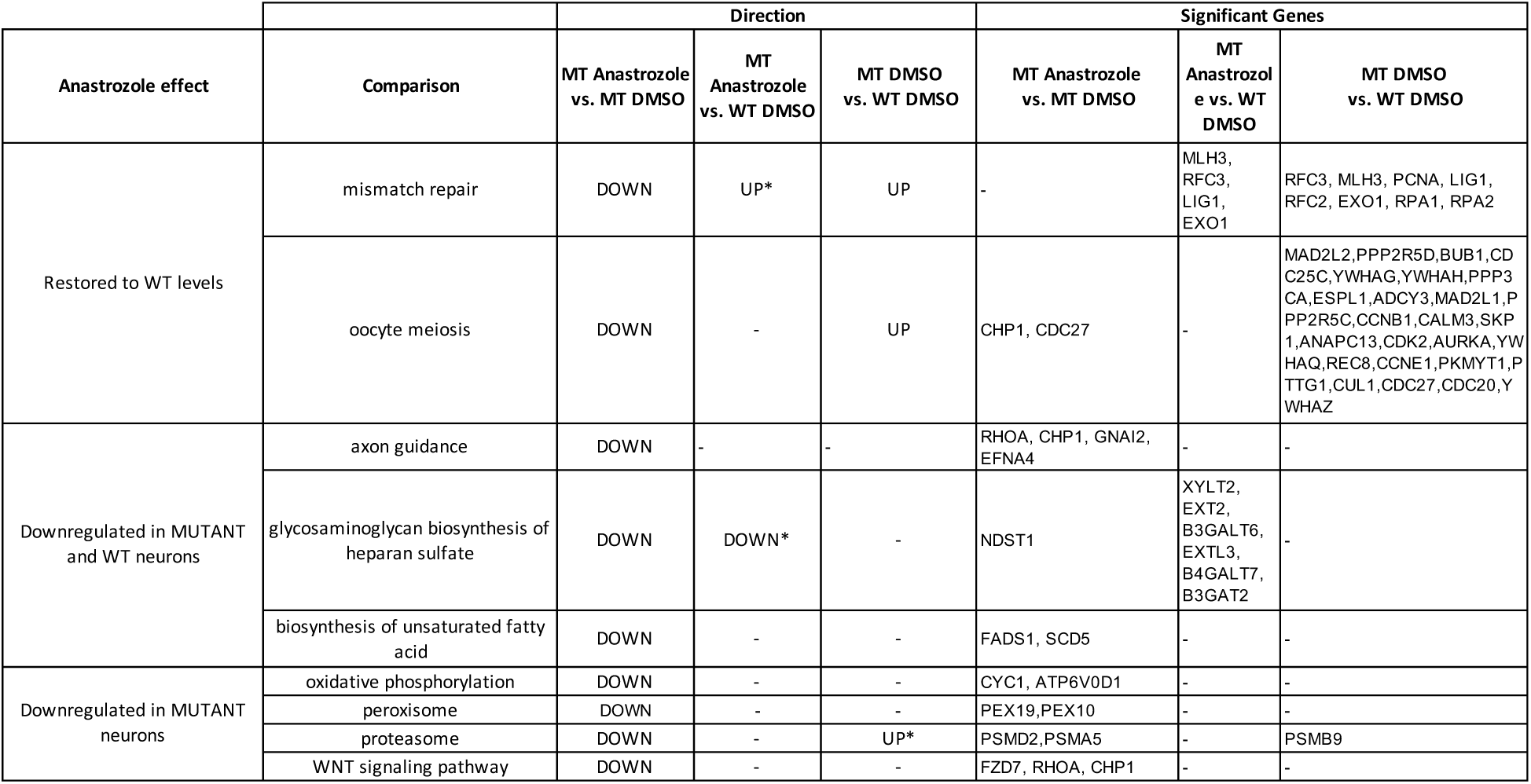
Selected significant pathways. MT, mutant *MA PT^R406W^* neuron. WT, wild-type Tau neuron. -, non significant q value for the comparison (> 0.25 FDR). *, q value (< 0.25). Significant genes with altered expression in each pathway.

Treatment with 10 ⍰µM anastrozole for 24 ⍰hours (MT Anast vs MT DMSO and WT Anast vs WT DMSO) led to significant downregulation of seven pathways—including axon guidance (NES = −1.86, q = 8.5×10^-3^), glycosaminoglycan biosynthesis of heparan sulfate (NES = −2.07, q < 1.5×10^-3^), and unsaturated fatty acid biosynthesis (NES = −1.96, q = 5.7×10^-^3) in both mutant and wild-type neurons (Fig. 5A, B, Suppl. Table 2,3), suggesting a reduction in surface heparan sulfate, a key mediator of Tau propagation ^27,28^. Anastrozole exerted a stronger effect on *MAPT^R406W^*neurons, suppressing 42 pathways such as oxidative phosphorylation (NES = −1.86, q = 8.6×10^-3^), proteasome (NES = −2.1, q = 1.2×10^-3^), Parkinson Disease (NES =-1.59, q = 3.8×10^-2^) and restoring oocyte meiosis to wild-type levels (n.s. MT Anast vs WT DMSO, Fig. 5A, B, Suppl. Table 4, 5). Analysis of the top 30 differentially expressed coding genes (Table 2) showed modest individual gene changes. While expression of key *MAPT^R406W^*-altered genes (*GRIN3A*, *COMT*, *NDNF*, *CHL1*, *NOVA1*, and *PQBP1*) remained unaffected, anastrozole upregulated genes such as *INTS6L*, *FBN3*, and *NPIPB12*, and downregulated *GRINA*, *DDOST*, and *FZD7*, known to play important roles in neuronal functions and associated with neurodegenerative disease ^29,30,31,32^.

**Table 2:**
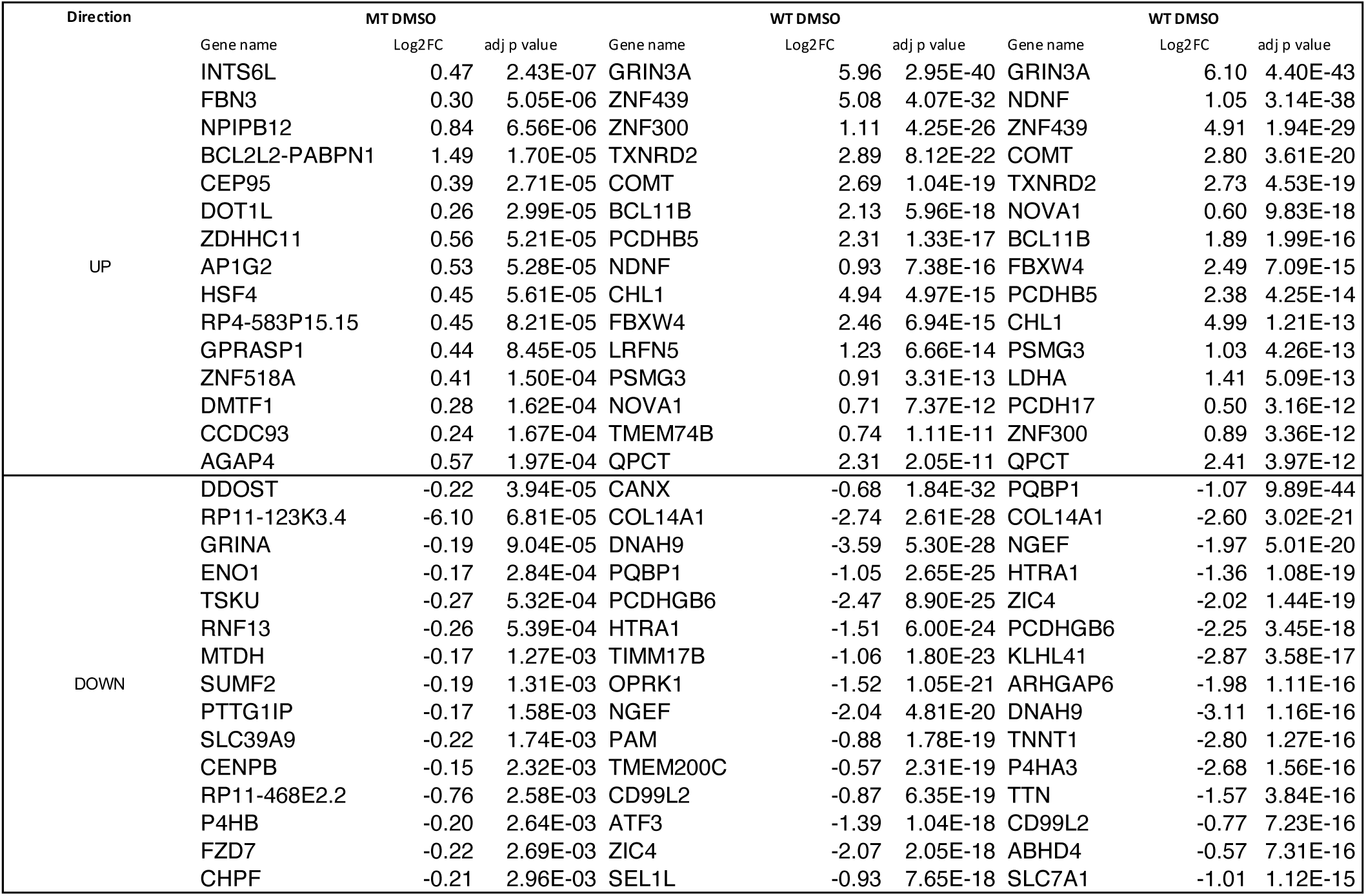
Top 30 significantly dysregulated genes across comparisons ranked on t-statistics.

Comparison of gene lists across treatment and genotype conditions (Fig. 5C, D, Suppl. Fig. 7C, D) led to identification of genes significantly dysregulated by mutation that were corrected by anastrozole treatment (Fig. 5E). Of interest, neuronal circuitry including *MUC6, DEF6*, and protocadherin A13 were restored to near WT levels by anastrozole. Overall, these drug-specific effects suggested a rebalancing of lipid homeostasis and oxidative phosphorylation pathways, possibly promoting neuronal health. The association with fatty acid levels and metabolism suggested potential novel targets for anastrozole such as the fatty acid metabolism regulator CYP4V2, a possibility we explored next.

### Drosophila screening identifies Cyp4 family as a candidate regulator of MAPT^R406W^ dysfunction

Having excluded estrogen and cholesterol pathways and identified new candidates as mediating anastrozole’s effects on *MAPT^R406W^*, we returned to the Drosophila model to identify and assess candidate targets. The DIOPT ortholog database identified several Drosophila *Cy p* genes as orthologs of human *CYP19A1/Aromatase* (Suppl. Table 6). *GMR*-driven RNAi knockdown of *Cyp4ac1*, *Cyp313a4*, or *Cyp316a1* each significantly rescued *GMR>hTau^R406W^*viability (Fig. 5F). Partial *Cyp4ac1* knockdown further enhanced rescue by low-dose (10 µM) anastrozole (Fig. 5G), indicating functional synergy. Although structurally similar to human CYP19A1/Aromatase, *Drosophila Cyp4ac1* is most orthologous with the human *CYP4* family, which plays an important role in lipid metabolism ^33^. Together, these findings suggest that anastrozole acts through a CYP4-like enzyme to rescue *GMR>hTau^R406W^*flies. Therefore, we investigated whether the neuroprotective effects of anastrozole could be attributed to its inhibition of CYP4V2.

### Knockdown and inhibition of CYP4V2 decreases ROS production in MAPT*^R406W^* lines

CYP4 enzymes are best known for their ω-hydroxylase function on arachidonic acid, transforming it into 20-hydroxyeicosatetraenoic acid (20-HETE). HET0016 is a small molecule that inhibits CYP4V- and CYP4Z-driven fatty acid oxidation activity ^34,35^. Treatment with 0.2 μM HET0016 strongly reduced rotenone-induced ROS production in *MAPT^R406W^* cells to levels approaching isogenic control cells using live imaging assay with CellRox Green (Fig. 6A, B). Further supporting a role for CYP4V2 in rescuing ROS production in *MAPT^R406W^* neurons, viral shRNA-mediated knockdown (KD) led to a strong decrease in rotenone-stimulated ROS production when compared to scrambled control shRNA (Fig. 6C, D). Live imaging with Cell Rox red was performed after green-tagged virus delivery and rotenone stimulation (Suppl. Fig. 8A, B). Lastly, we compared anastrozole (0.1 μM) and HETE0016 (0.1 μM) with a luminometry (RoxGlow) assay on cell lysate after 6 h treatment with rotenone, observing significant reduction of ROS production for both treatments using an independent method for ROS detection (Fig.6E). In order to ascertain whether CYP4V2 KD induced similar transcriptional changes as anastrozole treatment, we then examined the expression of MUC6. *CYP4V2* KD also significantly upregulated *MUC6*, indicating a shared mechanism of action between anastrozole and CYP4V2 targeting (Fig. 6F).

**Figure 6:**
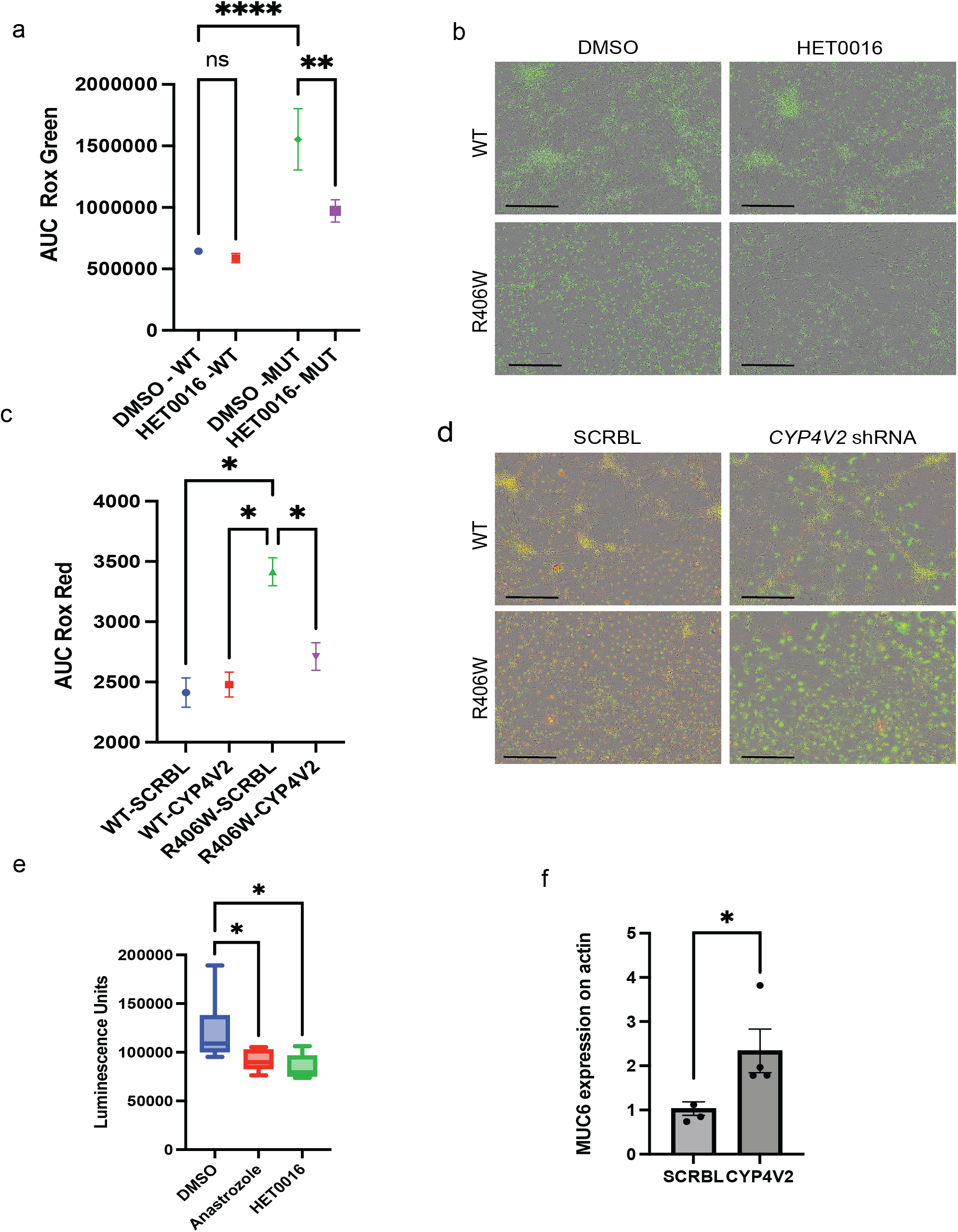
Chemical or genetic inhibition of CYP4V2 dampens ROS production in MAPT neurons. A) Quantification of ROS production with CellROXGreen (AUC of RCUxmm^2^), representative of two experiments with rotenone stimulation, comparing HET0016-treated cells versus control. B) Representative images of A at 25 h post-treatment (scale bars, 400 µm). C) Quantification of ROS production with CellROXRed (AUC of RCUxmm2), representative of two experiments with rotenone stimulation for scramble control shRNA (SCRBL)- or *CYP4V2* shRNA (CYP4V2)-treated neurons. D) Representative images of C at 25 h post-treatment (scale bars, 400 µm). \**p* < 0.05, \*\**p* < 0.01, and \*\*\*\**p* < 0.0001 in one way Anova with multiple corrections. ns, not significant. E) RoxGlow assay for quantification of reactive oxygen species in after 6 h of treatment in MAPT mutant neurons treated with 0.1 DM anastrozole, HET0016 in 0.1% DMSO control \**p* < 0.05, in one way Anova with multiple corrections. (F) qPCR for *MUC6* in SCRBL or *CYP4V2 MAPT*-mutant neurons, from two independent differentiations.

Together, these data support a role for CYP4V2 in *MAPT*-mediated ROS production and suggest a novel mechanism of action for anastrozole’s ability to protect *MAPT^R406W^* neurons.

## Discussion

In this study, we demonstrate the neuroprotective effects of the Aromatase inhibitor anastrozole in a *Drosophila hTau^R406W^* model and in a human tauopathy cell culture model. Using iPSC-derived cortical neurons from patients with the pathogenic *MAPT^R406W^* mutation, we demonstrated that anastrozole potently protected against oxidative stress independently of an effect on basal Tau phosphorylation. Surprisingly, we found that this neuronal protection was independent of anastrozole’s canonical role as an Aromatase inhibitor and was not dependent on its known off-target CYP27A. Instead, we propose CYP4V2 as a novel target of anastrozole that mediates its rescue of both Drosophila and human cortical neuron models.

Basal Tau phosphorylation is essential for normal development, but pathological stressors—including mitochondrial dysfunction, inflammation, amyloid pathology, and *MAPT* mutations—can drive aberrant phosphorylation via kinase activation and phosphatase inhibition ^36,37,38^. This leads to Tau misfolding, aggregation, and a feed-forward cycle of mitochondrial and lysosomal dysfunction with oxidative damage ^39,40^. Unlike prior *MAPT* iPSC studies ^4,10^, we did not detect Tau hyperphosphorylation in *MAPT*^R406W^ neurons, likely due to early differentiation (4 weeks) and the specific effect of the *R406W* mutation, which impairs microtubule binding and causes mislocalization without necessarily altering phosphorylation ^41^. While hyperphosphorylation was absent, an anastrozole-induced reduction in basal Tau phosphorylation may be protective in models with pathological phosphorylation.

Although Aromatase inhibitors have been associated with adverse cognitive effects in clinical settings ^42^, our findings indicate that anastrozole confers neuroprotection through a mechanism independent of Aromatase inhibition, estrogen depletion, or cholesterol sterol oxidation. Neither an alternative Aromatase inhibitor nor an estrogen receptor antagonist replicated anastrozole’s effects, and no reduction in estrogen levels was detected following treatment, indicating a mode of action distinct from its known activities. In *Drosophila*, which lack a functional aromatase ortholog, the rescue of tauopathy phenotypes by anastrozole was linked to Cyp4ac1, suggesting that its neuroprotective activity involves targeting a different CYP family member other than CYP19A1. This was further supported by our iPSC-derived neuron studies, which showed that anastrozole had no effect on 27-hydroxycholesterol (27-OHC) levels, a proposed off-target ^43^. Together, these data position anastrozole as a candidate neuroprotective agent acting via a novel CYP4-dependent pathway.

As the target of anastrozole in human iPSC-neurons was unknown, we carried out transcriptomic analyses to identify functional pathways that may highlight candidate target genes. Gene set enrichment analyses showed anastrozole corrected pathways altered in mutant neurons, such as oocyte meiosis, and suppressed oxidative phosphorylation, a driver of oxidative stress and neuronal death, thereby supporting our functional data. Furthermore, these data led us to speculate that anastrozole may alter lipid metabolism and support a resilience phenotype by decreasing oxidative phosphorylation, with potential additional beneficial effects on prion-like Tau spread due to its downregulation of heparan sulfate-related pathways ^27,28^.

Transcriptomic analysis of iPSC neurons, combined with ortholog screening in Drosophila, identified CYP4V2 as a likely off-target of anastrozole. Inhibition of *CYP4V2* expression reduced ROS production in our models, supporting its potential role in mediating the neuroprotective effects of anastrozole. As a ω-hydroxylase of arachidonic acid, CYP4V2 generates 20-HETE, a byproduct detrimental for neuronal health ^44^. These data also show that anastrozole restored expression of *PCDHA13* and *MUC6* — key regulators of neuronal adhesion ^45^, which were downregulated in *MAPT*-mutant neurons. *MUC6* was also upregulated by *CYP4V2* KD in the same model, supporting this as one of the possible targets downstream of CYP4V2 loss. Prior studies implicate CYP4 activity and 20-HETE in ROS amplification via NADPH oxidase ^35^. HET0016, a CYP4 inhibitor, similarly reduced ROS and pyroptosis in traumatic brain injury models ^44,46,47^. We confirmed that HET0016 suppressed ROS in *MAPT^R406W^* neurons at low micromolar concentrations, consistent with CYP4V2 inhibition as mechanistic contributor to the protective effects of anastrozole and a potential therapeutic target in *MAPT*-associated tauopathies. The robust reduction of oxidative stress in neurons with anastrozole, CYP4V2 inhibitor and KD was detected with two independent methods: observing increase in fluorescence over time or by luminescence endpoint assay, suggesting that suppression of intracellular reactive oxygen species supports prolonged neuronal survival to stress exposure.

While our work indicates that CYP4V2 is a promising therapeutic target for tauopathy, it is important to note that homozygous loss of function polymorphisms within *CYP4V2* reduce fatty acid hydroxylation ^34^ and are linked to Bietti crystalline dystrophy (BCD), a disease that can lead to impaired vision ^48^. This suggests that additional research into the proper targeting of the CYP4V2 network will be required when considering therapeutics. Although fly models are valuable for drug screening ^49–54^, differences in orthologs between species might also explain the additive—not synergistic—effect of low-dose anastrozole in the *Cyp4ac1* KD model. Possible compensatory effects of other orthologs cannot be ruled out, but the *Drosophila* model provided a valuable screening tool for drug identification and subsequent target prioritization before evaluation in mammalian models.

In summary, we have demonstrated that anastrozole confers potent protection against oxidative stress in a human neuronal model of tauopathy. We provide evidence that this protection is through an Aromatase- and estrogen-independent mechanism; we propose that this neuroprotective effect may be mediated through CYP4V2 inhibition. However, anastrozole does not cross the blood-brain barrier and Aromatase inhibition may impair cognitive function ^13^; furthermore, anastrozole is metabolized to hydroxy-anastrozole by CYP3A4 *in viv*^5^*o*^5^, which may alter its binding affinity for CYP4V2. Direct binding studies would be beneficial to identify key functional groups and interacting motifs with CYP4V2, informing rational design of optimized lead compounds that more precisely target CYP4V2 to treat tauopathies including Alzheimer’s and related dementias.

## Methods

### Fly stocks

*UAS*-*hTau*^WT^ and *UAS*-*hTau^R406W^* flies ^2^ were crossed with *GMR*-*gal4* to generate *GM R*>*hTau*^WT^ and *GMR*>*hTau^R406W^* flies, respectively, as *Drosophila* tauopathy models. Human orthologs of fly genes were predicted by *Drosophila* RNAi Screening Center (DRSC) integrative ortholog predictive tool DIOPT ^56^. Fly stocks carrying small interfering RNA (siRNA) for *Cyp4ac1*, *Cyp313a4*, or *Cyp316a1* were obtained from Bloomington *Drosoph ila*Stock Center (BDSC) and crossed with *GM R*>*hTau^R406W^*flies to generate a partial knockdown of these genes at 23 °C.

### iPSC derived neurons

iPSCs were differentiated into neural progenitor cells (NPCs) as previously described ^57^. NPC identity was confirmed by visualization of the expression of common NPC markers Nestin, SOX2, PAX6 and FOXP2 (Suppl. Fig. 1B). NPCs were differentiated to forebrain neuronal cultures with a 4 week long differentiation as previously described ^8^, and neuronal identity was confirmed by expression of Tau, TUJ1 and MAP2 after 4 weeks of differentiation (Suppl. Fig. 1C). NPC were provided by Neuracell ^58^, official distributor of iPSC generated by the Tau consortium ^7^

All NPC were from female donos. The Isogenic set 11362 (R406W) and 11362-C11 (corrected WT) was APOE 23 carrier, while 11421 (R406W) and 11421-2A07 (corrected WT) was APOE 33 carrier.

### *Drosophil*a chemical testing

Vismodegib, bortezomib, anastrozole, JQ1, and idelalisib (Selleck Chemicals, Houston, TX) were dissolved in DMSO (Sigma Aldrich; St. Louis, MO) to prepare stock solutions. Fly food with chemicals (0.1% final DMSO concentration) was aliquoted into plastic vials (Thermo Fisher Scientific; Waltham, MA), and *GMR*>*hTau^R406W^* progenies were cultured until adulthood on fly medium for 17 days at 18 °C. The number of adults was divided by that of total pupae to calculate percent viability.

### Stress susceptibility-rotenone

For oxidative stress susceptibility assays, Incucyte live-cell imaging was used on four-week-old neurons treated with rotenone (500 nM, 1 µM, or 3 µM; Sigma Aldrich) or DMSO, and were monitored for 24 hours with images acquired every hour. Cells were pre-treated for 24 hours with the compound of interest, then treated with the described concentration of rotenone and CellRox Green (1:500). Oxidative stress was quantified by measuring the integrated intensity of CellRox Green fluorescence (1:500; Thermofisher Scientific) normalized to cell confluency. For the preliminary screen of the initial compounds to rescue rotenone-induced stress susceptibility (Suppl. Fig 2B), neurons were first gently dissociated with accutase (ICT, AT-104) and re-seeded prior to treatment, resulting in a more sensitive stress response. For these experiments, oxidative stress was assayed over the course of 1 hour, with images taken every 5 minutes. Compounds were diluted in DMSO 1 M-10 µM to a final 1% DMSO. Vismodegib, bortezomib, anastrozole, JQ1, idelalisib, rapamycin, exemestane (steroidal Aromatase inhibitor; Selleck), fulvestrant (Estrogen receptor antagonist; Selleck), bicalutamide (27-OHC inhibitor; Selleck), efavirenz (24-OHC inhibitor; Sigma Aldrich), and fluvoxamine (27-OHC activator; Sigma Aldrich) were provided from the Cagan’s Lab. HET0016 was purchased from MedChemExpress (HY-124527).

### Immunofluorescence

NPCs were seeded at 2×10^5/well in 24-well plates, and were fixed with Formalin (Sigma-Aldrich) for 15 minutes at room temperature prior to washing with PBS, permeabilization in 0.1% Triton-×100 (Thermo Fisher Scientific) and blocking in 1% BSA in PBS. Cells were labelled for Nestin (Abcam ab22035 1:100), SOX2 (NEB 3579S 1:400), PAX6 (Abcam ab5790 1:100) and FOXP2 (Abcam ab16046 1:100) antibodies overnight at 4C to evaluate neuronal progenitor multipotency. Alexa-fluor secondary antibodies (Thermo Fisher Scientific) were incubated at 1:100 for 2 hours at room temperature, followed by 10 minutes incubation with 1 ug/ml DAPI. Cells were imaged on a Leica DMIL LED Inverted Routine Fluorescence Microscope with a 20x objectiveFor neuronal characterization NPC were seeded in a 8 chamber slide (CELLTREAT 229168) at 2×10^4^/mL, 250 uL/chamber. After 4 weeks of differentiation cultures were fixed with 4% PFA (Thermo Fisher Scientific) in PBS for 10 min, permeabilized with 0.1% Triton-×100 (Thermo Fisher Scientific) in PBS for 20 min and blocked with 2% BSA in PBS for 1 h (Miltenyi Biotech 130-091-376). Staining was performed in blocking buffer for MAP2 (Abcam ab5392 1:50,000), NeuN (Abcam ab104225 1:1,000), tubulin beta 3 (BioLegend mms-435p 1:500) O/N at 4 °C. Secondary antibodies were incubated 1:100 in blocking buffer for 2 h at RT. Images were acquired with Keyence Microscope at 10x magnification after mounting with Prolong antifade containing DAPI (Sigma Aldrich).

### ELISA and biomolecule detection

We measured the phosphorylation of three Tau residues in *MAPT^R406W^* and WT neurons by harvesting cell lysates for ELISA after 24 h treatment with small molecules. We detected phosphorylated serine 199 (S199, Thermo Fisher Scientific KHB7041), serine 396 (S396, (Thermo Fisher Scientific KHB7031) and threonine 181 (T181, Thermo Fisher Scientific KHO0631). Relative levels of phosphorylation between genotypes were calculated by normalising pTau absorbance values to total Tau absorbance from the same lysate (Thermo Fisher Scientific KHB0041). To assess the effect of small molecules on tau levels, the absorbance for baseline DMSO condition was subtracted from the absorbance measurements of each treatment condition from the same iPSC line. LDH assay was performed with Promega kit (G1780). Briefly, Neurons were treated with 1 nM to 10 µM of each compound for 24 h and supernatant was harvested for assay. E2 estradiol (Thermo Fisher Scientific KAQ0621) and testosterone (Abcam ab108666) were detected in supernatants of neurons treated with 10 nM to 10 µM of each compound for 24h. 24-hydroxycholesterol (Enzo Life Sciences ADI-900-210-0001), 27-hydroxycholesterol (MyBioSource MBS7606791) and total cholesterol (Amplex™ Red Cholesterol Assay Kit A12216) quantification was performed on supernatant after 24 h from treatment with selected compounds. RoxGlow assay to detect ROS production in neurons was performed according to the manufacturer’s instruction with a 6 h incubation with rotenone before detection.

### *CYP4V2* targeted knockdown

The viral vector designed by Vectorbuilder was able to induce *CYP4V2* shRNA expression in mature neurons under the U6 promoter (Suppl. Fig. 1D) to avoid developmental effects of CYP4V2 loss in neurons. Briefly, after 24 h incubation with virus of MOI 10 the neuronal media was replaced and incubated for an additional 48 h before initiating the assay. Expression of GFP was used as a readout for evaluation of efficient transfection. To validate efficient KD of *CYP4V2*, we extracted RNA and proteins from shRNA-treated neurons and performed RT-qPCR for *CYP4V2* with Taqman gene expression assay (Thermofisher Hs01389878_m1) performed with fast advanced mastermix (Thermo Scientific 4444557) (Suppl. Fig.1E). Data is representative of two independent differentiations and transfections on one iPSC isogenic set, with representative images shown in (Suppl. Fig. 8B). QPCR for *MUC6* was performed using the primers (FWD 5’-GGACTGTGAGTGTCTGTGCGAT-3’, REV 5’-GCGTGTTGTAGAAGCCGCAGTA-3’) with the SYBR Green master mix (Thermo Scientific 4385618).

### RNA-seq analysis

RNA was isolated from the neurons (RNeasy Mini, Qiagen) and sequencing (2×150 bp) was performed by Genewiz. Reads were processed using STAR (Spliced Transcripts Alignment to a Reference) for alignment to the GRCh38 reference genome (Gencode v38 annotation). The STAR index was pre-built using Gencode v38 to optimize splice-aware alignment. Sequencing reads were aligned with STAR using default parameters with additional specifications for handling multi-mapping reads and retaining unaligned reads for quality assessment. Each sample’s alignment process generated a sorted BAM file as output, which served as input for further downstream quantification. Following alignment, transcript abundance was estimated using RSEM (RNA-Seq by Expectation Maximization). RSEM was run with paired-end mode enabled, using the same Gencode v38 annotation to ensure consistency with the STAR alignment step. Gene-level and transcript-level expression estimates were obtained, and gene expression counts were summarized for downstream normalization and analysis. After quantification, quality control (QC) metrics were collected for each sample using Picard tools. Picard’s CollectAlignmentSummaryMetrics, CollectRnaSeqMetrics, and CollectGcBiasMetrics were used to assess total read counts, duplication rates, ribosomal RNA contamination, strand specificity, intergenic and intronic base proportions, and GC bias. Additional duplicate removal was performed using Picard’s MarkDuplicates to correct for technical biases introduced during library preparation and sequencing. These QC metrics were extracted from Picard output files and compiled into a structured dataset for review. Following quality assessment, sample-level sequencing metrics were summarized using a Python script that parsed the Picard QC outputs to extract key values such as percentage of ribosomal RNA, total aligned reads, read strand balance, and duplication rates. This allowed for systematic identification of outlier samples with low-quality data.

Expression data underwent count-per-million (CPM) filtering to remove lowly expressed genes. Genes were retained if they had CPM > 0.3 in at least 30% of samples, ensuring inclusion of biologically relevant transcripts. Genes with effective transcript lengths ≤ 50 bp were also removed to prevent artifacts in subsequent analyses. Conditional quantile normalization (CQN) was applied to correct for GC content and gene length biases, followed by limma-trend normalization to account for remaining technical variation. Principal component analysis (PCA) was performed on the normalized expression data to visualize sample clustering, detect potential batch effects, and identify outliers. The number of principal components retained varied by dataset and was determined based on the proportion of variance explained. PCA plots were generated for each dataset, with coloring based on relevant covariates such as haplotype, ancestry, and batch effects to ensure appropriate normalization and batch correction. Differential gene expression analyses were conducted using DESeq2, incorporating relevant covariates tailored to each dataset to control for biological and technical confounders.

Differential gene expression results were visualized using volcano plots to highlight significantly upregulated and downregulated genes. For each comparison, genes were plotted based on their log2 fold change and statistical significance (-log10 *p*-value or -log10 adjusted *p*-value), with significance thresholds set at α = 0.05. Genes meeting significance criteria were colored accordingly, with upregulated genes in red and downregulated genes in blue in the volcano plots.

To further explore the biological relevance of differentially expressed genes, gene set enrichment analysis (GSEA) was performed using pre-ranked lists generated from DESeq2 results. Genes were ranked based on their test statistic (Wald statistic) ^59^. Genes with missing values (or NA) in key columns (*p*-value, adjusted *p*-value (padj), log2 fold change, or test statistic) were removed before ranking. Enrichment analysis was conducted separately for each comparison using curated gene sets from Gene Ontology (GO) biological processes, KEGG pathways, and Reactome pathways. Pathways were considered statistically significant based on a false discovery rate (FDR) threshold, with results filtered at multiple significance levels. The normalized enrichment score (NES) was calculated to quantify the strength of pathway enrichment.

To compare pathway enrichment patterns across multiple tissues and analyses, shared and unique pathways were identified through hierarchical clustering and heatmap visualization. The significance of each pathway in each dataset was denoted using asterisks (***FDR ≤ 0.01, **FDR ≤ 0.05, *FDR ≤ 0.25). In Python, values smaller than approximately 10^-308^ underflow to zero due to floating-point precision limits. To account for this, extremely small Q-values from pathway analysis were reported as 1.00^-10^ where applicable. Raw FASTQ RNA-seq data and count files are deposited in GEO: GSE294023.

## Supplementary Tables

**Suppl. Table 1:** FDA approved Cancer drug tested on Drosophila model.

**Suppl. Table 2:** RNA-seq DEG WT-Anast vs WT-DMSO and pathway analysis.

**Suppl. Table 3:** RNA-seq DEG MT-Anast vs MT-DMSO and pathway analysis.

**Suppl. Table 4:** RNA-seq DEG MT-DMSO vs WT-DMSO and pathway analysis.

**Suppl. Table 5:** RNA-seq DEG MT-Anast vs WT-DMSO and pathway analysis.

**Suppl. Table 6:** Orthologs and analogs of CYP between Drosophila and Human.

## Supplementary Figures

**Suppl. Fig. 1:** Characterization of iPSC lines used in the study and functional data.

**Suppl. Fig. 2:** Screening of six compounds in MAPT*^R406W^* neurons reveals differential abilities to reduce rotenone-induced ROS production.

**Suppl. Fig. 3:** Alternative Aromatase inhibitors have no effect on ROS production or Tau phosphorylation.

**Suppl. Fig.4:** Anastrozole does not affect E2 estradiol or testosterone levels in vitro.

**Suppl. Fig.5:** Cholesterol biosynthesis is altered in MAPT*^R406W^* neurons, but targeting does not recapitulate anastrozole treatment.

**Suppl. Fig. 6:** Hydroxycholesterol production in MAPT*^R406W^* neurons and isogenic controls.

**Suppl. Fig. 7:** Additional data from RNA-seq analysis.

**Suppl. Fig. 8:** Validation of CYP4V2 knockdown.

## Acknowledgements/Conflicts/Funding Sources

We thank Dr. Celeste Karch for providing iPSC lines generated by the Washington University facility, and Dr. Mel B. Feany for providing *hTau* transgenic flies. This work was supported by funding from the Rainwater Charitable Foundation (AMG, KRB, RLC). UK Dementia Research Institute (UKDRI-4211) through UK DRI Ltd, principally funded by the UK Medical Research Council (KRB). RC, MS, KB, and AG have filed a patent on use of small molecule anastrozole analogs inhibiting CYP4V2 to reduce ROS production in neurodegenerative diseases.

## Authors contribution

RC, CP, SAW, and KRB were responsible for writing the initial draft. MS was responsible for the studies in *Drosop hil a*. KRB and DAP were responsible for drug screening in ROS assays and ELISAs. CP and SAW were responsible for RNA-seq analysis. CP and JO were responsible for *CYP4V2* KD studies. All the authors have reviewed and approved the final manuscript.

---

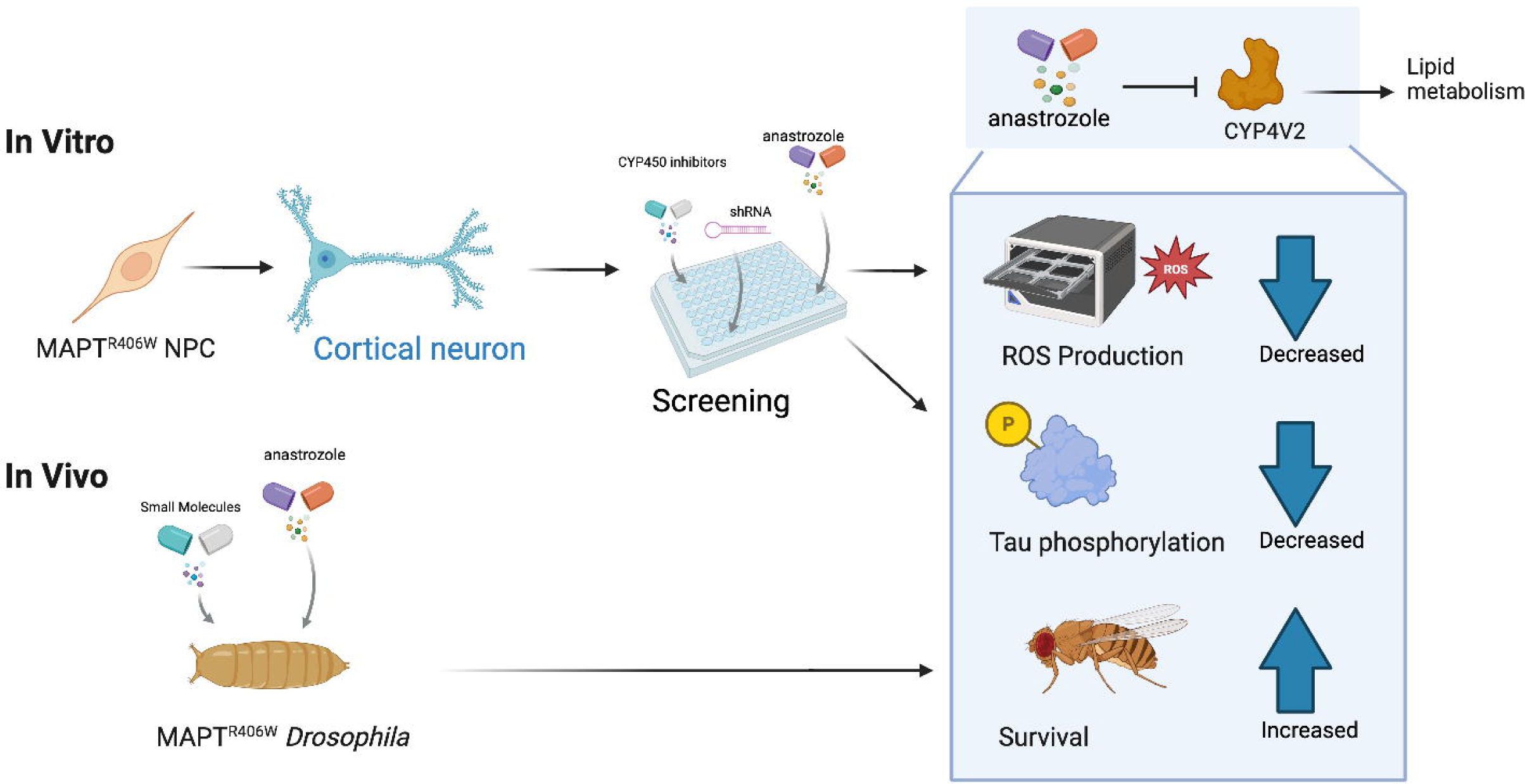

