## Supplementary Figures for "Alternative Anastrozole Target CYP4V2 Impairs Neurons in Preclinical Models of Inherited Tauopathy"

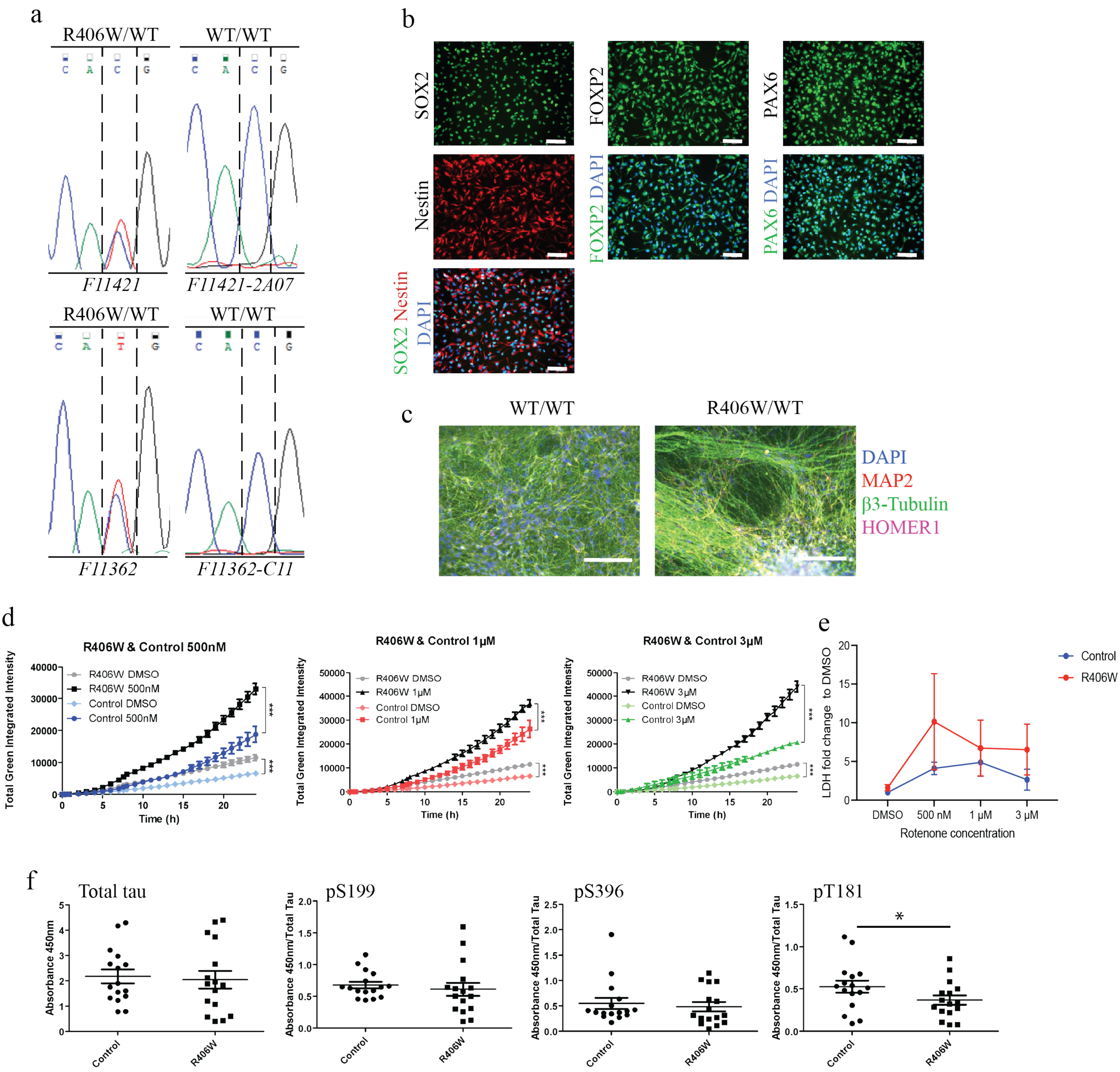


**Suppl. Fig. 1: Characterization of iPSC lines used in the study and functional data.** A) Genotype traces for F11421 and F11362 isogenic sets. B) NPC characterization by immunofluorescence shows expression of neural progenitor markers: SOX2, FOXP2, NESTIN, and PAX6, with DAPI to identify nuclei. Scale bars = 100 µm. C) Validation of differentiated neurons with MAP2, Homer1 and beta 3 tubulin, with DAPI. Scale bars, 150 µm. D) Quantification of CellROX Green in control and mutant cells with rotenone or unstimulated (DMSO). Representative of one experiment on a single isogenic set, averaged across three wells each with three fields of view. Linear regression, $*p < 0.05$, $**p < 0.01$, and $***p < 0.001$. Error bars = SEM E) Extracellular release of LDH after 24 h treatment (DMSO or rotenone), representative of two experiments on two isogenic sets, two-way Anova statistical test was used to compare the groups. F) Total Tau and phosphorylated Tau in 4 weeks old $MAPT^{R406W}$ neurons versus control representative of four independent experiments in two isogenic sets, averaged over two technical replicates per condition. Student's t-test was used.

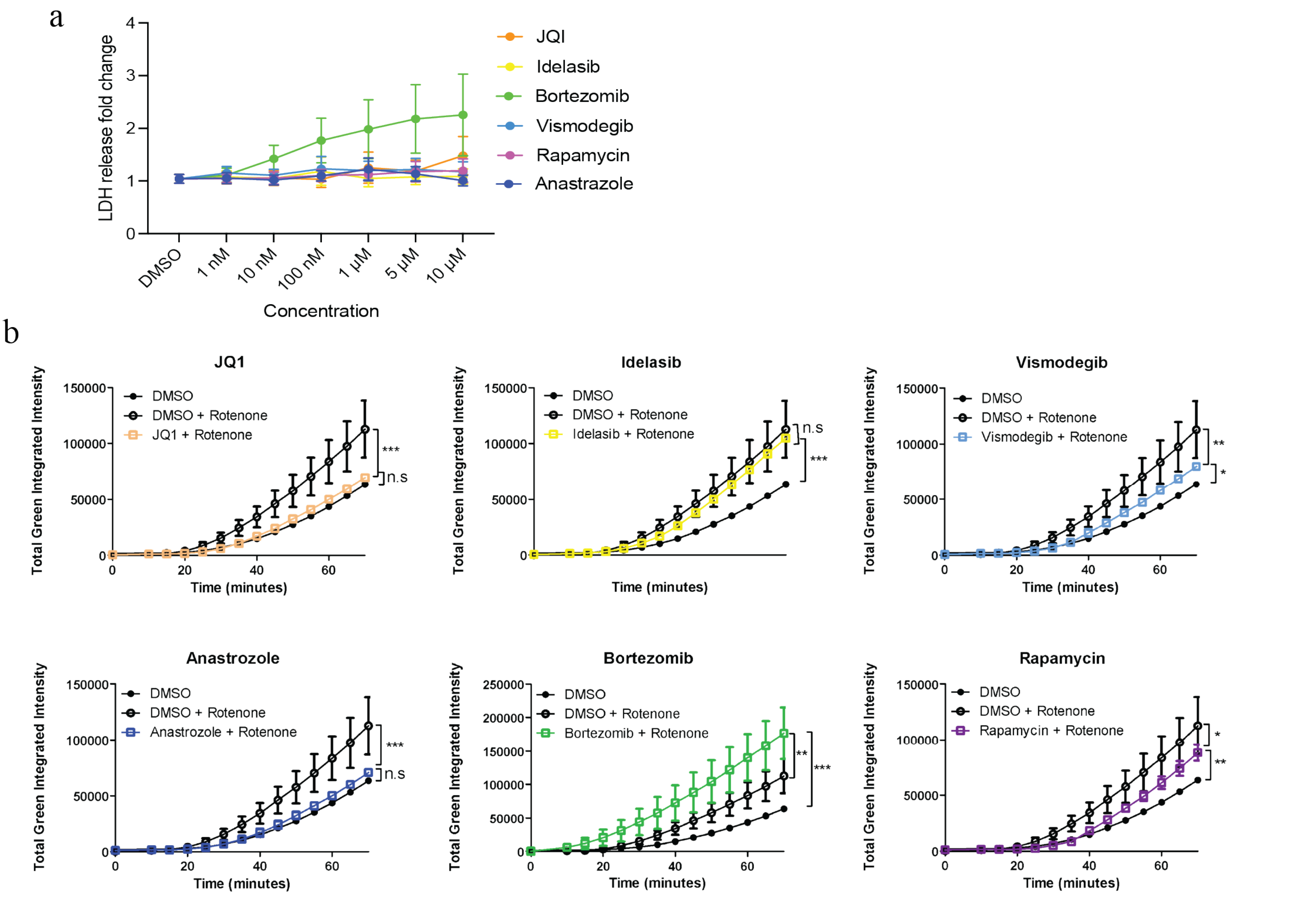


**Suppl. Fig. 2: Screening of six compounds in MAPT$^{R406W}$ neurons reveals differential abilities to reduce rotenone-induced ROS production.** A) LDH assay on 6 selected compounds incubated with iPSC-neurons for 24 hours at concentrations of 1 nM to 10 µM. Data normalized to LDH levels in DMSO control. Representative of three experiments incorporating 1 isogenic set of *MAPT*$^{R406W}$ neurons (N = 2 x *MAPT*$^{R406W}$, 1 x WT), statistics performed with two way Anova. B) Quantification of CellRox fluorescence intensity upon rotenone stimulation for these compounds at 1 µM. Data points represent an average of three technical replicates, each consisting of three fields of view, derived from a single experiment conducted in one*MAPT*$^{R406W}$ line. Linear regression, **$p < 0.05$, **$p < 0.01$, and ***$p < 0.001$. n.s, not significant.

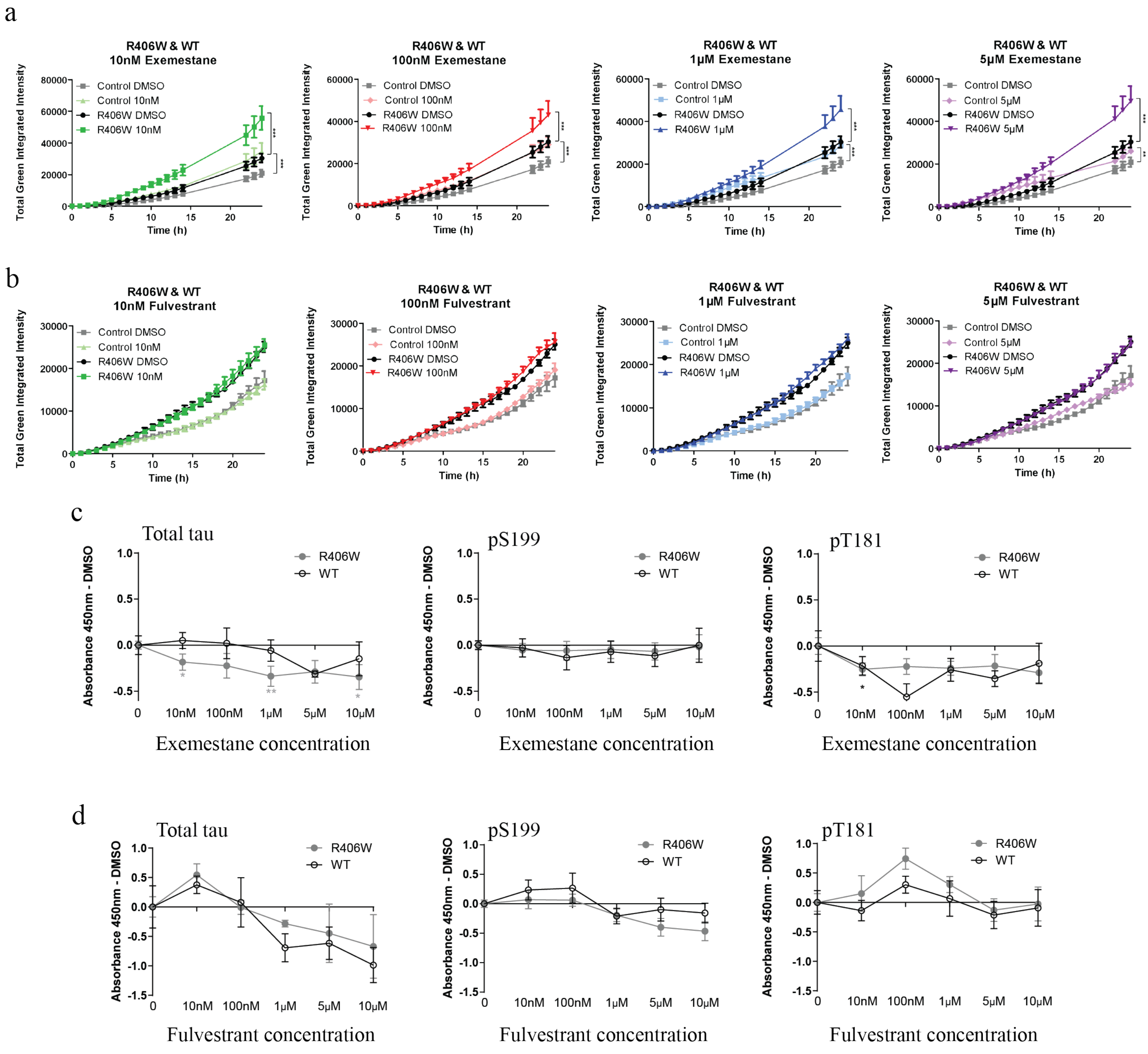


**Suppl. Fig. 3: Alternative Aromatase inhibitors have no effect on ROS production or Tau phosphorylation.** A-B) CellROX green fluorescence intensity in *MAPT*$^{R406W}$ neurons and controls following treatment with exemestane (A) or fulvestrant (B) prior to exposure to rotenone. C-D) Relative total Tau and phospho-Tau levels following 24 h exposure to either exemestane (C) or fulvestrant (D), compared to untreated DMSO controls. Data points represent N = 2 independent isogenic pairs, each consisting of three technical replicates encompassing three fields of view. A-B assessed by linear regression, C-D by two-way ANOVA with Dunnett's multiple comparisons test. Error bars = SEM. *p < 0.05, **p < 0.01, and ***p < 0.001.

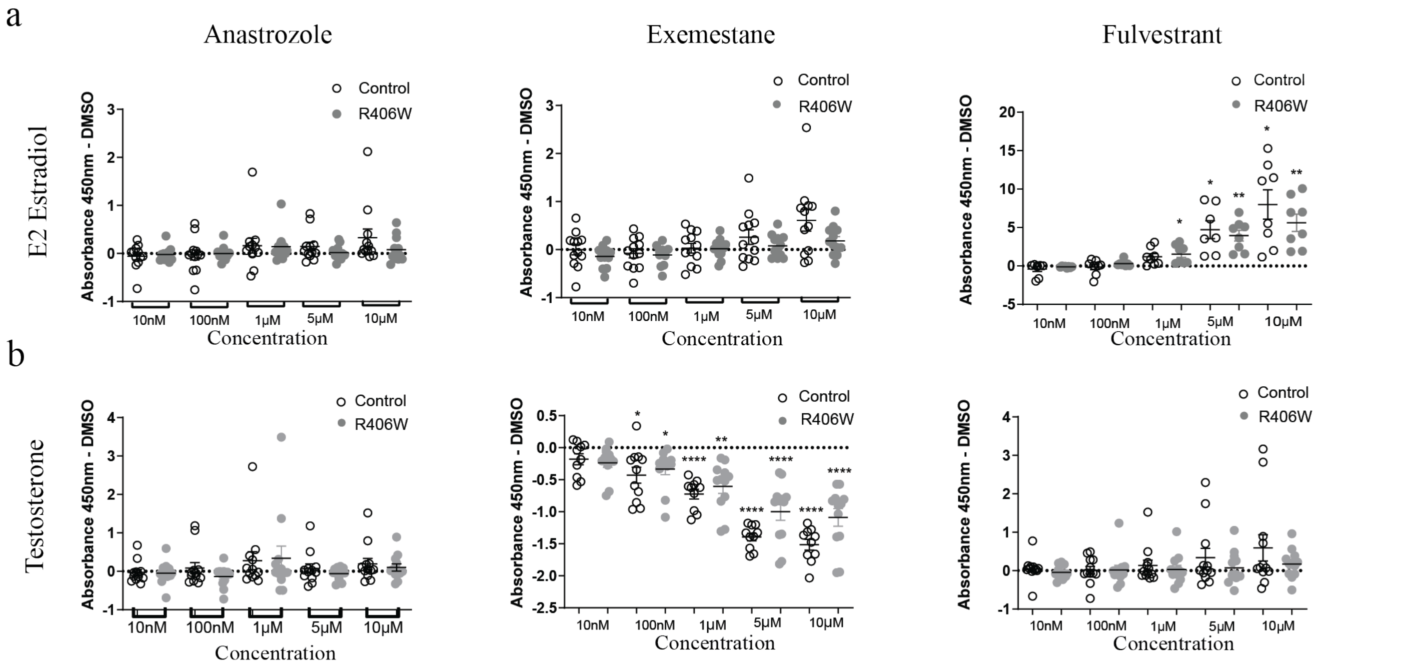


**Suppl. Fig.4: Anastrozole does not affect E2 estradiol or testosterone levels in vitro.** A-B) Relative extracellular E2 estradiol (A) and testosterone (B) levels from *MAPT*$^{R406W}$ neurons and isogenic controls following 24 h treatment with anastrozole, exemestane or fulvestrant, compared to untreated DMSO. Representative of two independent experiments on two isogenic sets, averaged from 3 technical replicates per condition. Two-way ANOVA with Dunnett's multiple comparisons test. Error bars = SEM. $*p < 0.05$, $**p < 0.01$, and $****p < 0.0001$.
.

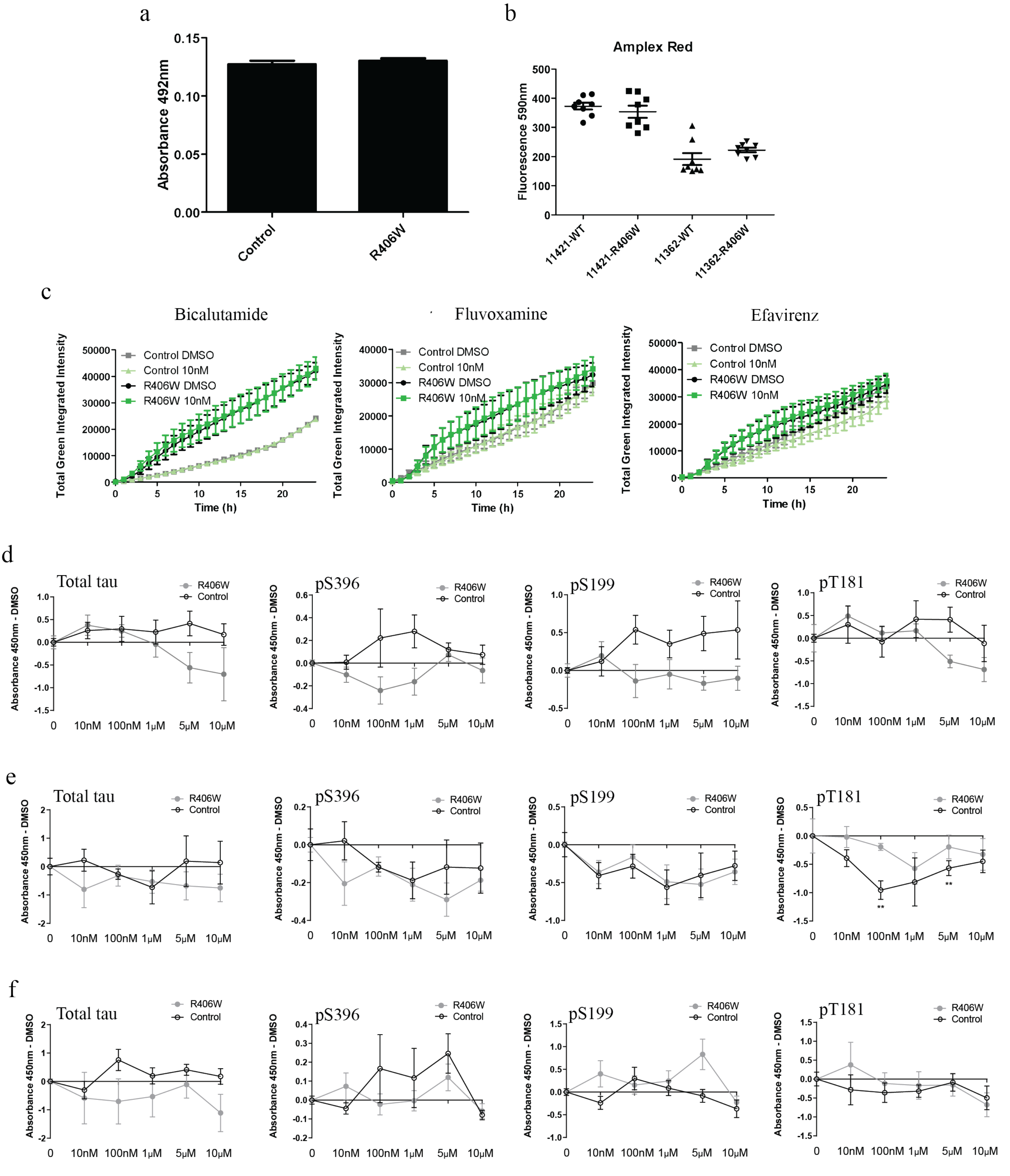


**Suppl. Fig.5 : Cholesterol biosynthesis is altered in $MAPT^{R406W}$ neurons, but targeting does not recapitulate anastrozole treatment.** *A) Quantification of neutral triglycerides by Oil Red O staining. B) Quantification of total cholesterol with Amplex RedTotal lipid in control (WT) and $MAPT^{R406W}$ neurons, split by donor iPSC line. Representative of a single experiment in two isogenic sets, averaged over 4 technical replicates, Student's t-test statistic test used. C) CellROX Green fluorescence intensity over time in control and $MAPT^{R406W}$ neurons treated with 10 nM bicalutamide, fluvoxamine, or efavirenz for 24 h prior to rotenone exposure. Data points represent N = 2 independent isogenic pairs, each consisting of three technical replicates encompassing three fields of view, linear regression. Error bars = SEM. D-F) Relative levels of total Tau and phospho-Tau in control and $MAPT^{R406W}$ neurons following 24 hours of treatment with bicalutamide (D), fluvoxamine (E), or efavirenz (F). Representative of two independent experiments on two isogenic sets, averaged from 3 technical replicates per condition. Two-way ANOVA with Dunnett's multiple comparisons test. Error bars = SEM. **p < 0.01.*

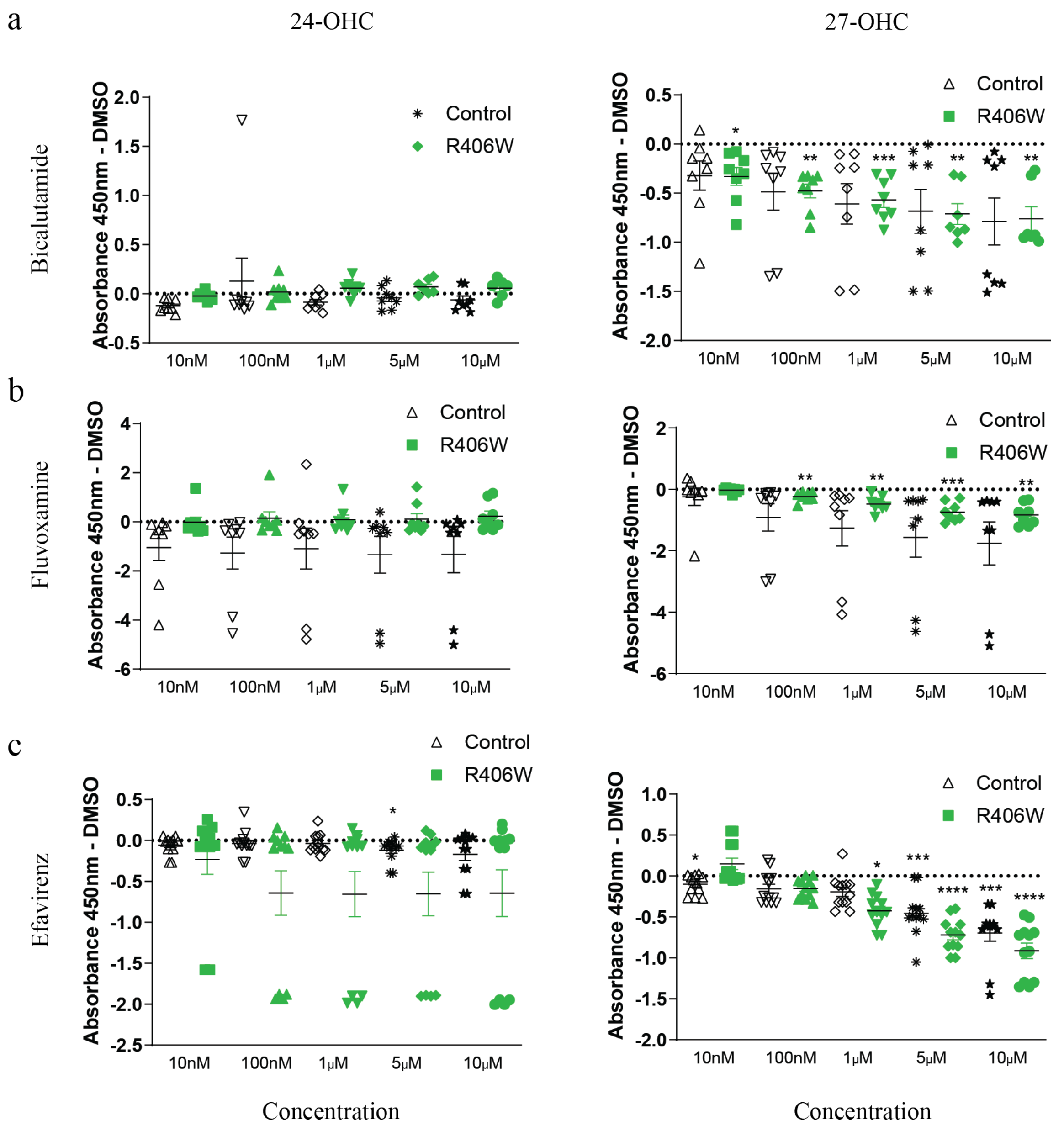


**Suppl. Fig. 6: Hydroxycholesterol production in MAPT$^{R406W}$ neurons and isogenic controls.** Production of 24-OHC (left) or 27-OHC (right) upon 24 h stimulation with A) bicalutamide, B) fluvoxamine, and C) efavirenz, normalised to DMSO control. Representative of two independent experiments on two isogenic sets, averaged from 3 technical replicates per condition. Two-way ANOVA with Dunnett's multiple comparisons test. Error bars = SEM. *$p < 0.05$, **$p < 0.01$, ***$p < 0.001$, and ****$p < 0.0001$ .

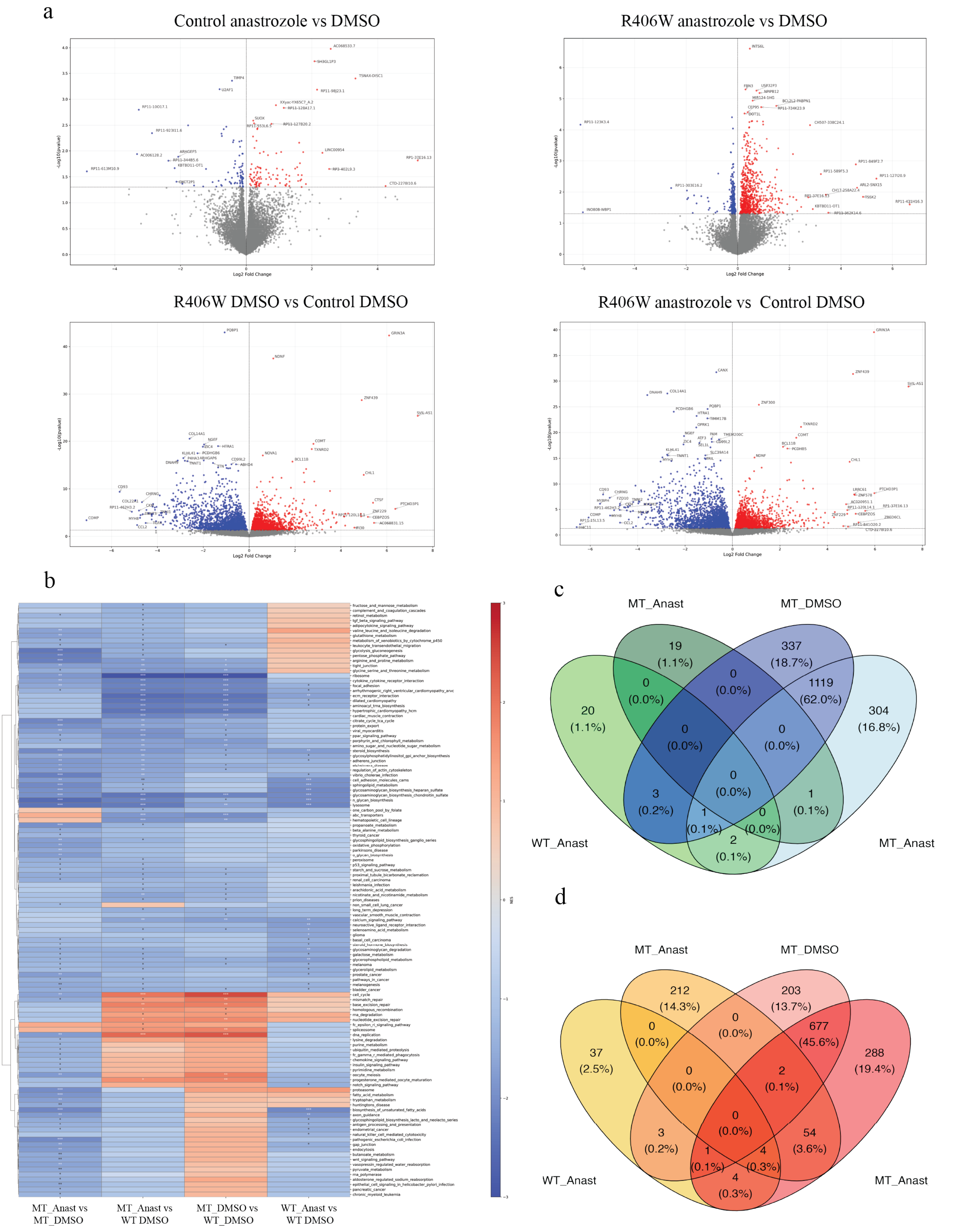


**Suppl. Fig. 7: Additional data from RNA seq analysis.** A) Volcano plots showing differential gene expression between *MAPT*[R406W] and control iPSC neurons with and without anastrozole treatment. Red and blue dots represent significantly upregulated and downregulated genes, respectively (p-value < 0.05, |log2FoldChange| > 0). B) GSEA pathway enrichment (NES) from all 4 comparisons. C-D) Overlapping down (C) and upregulated (D) genes across conditions.

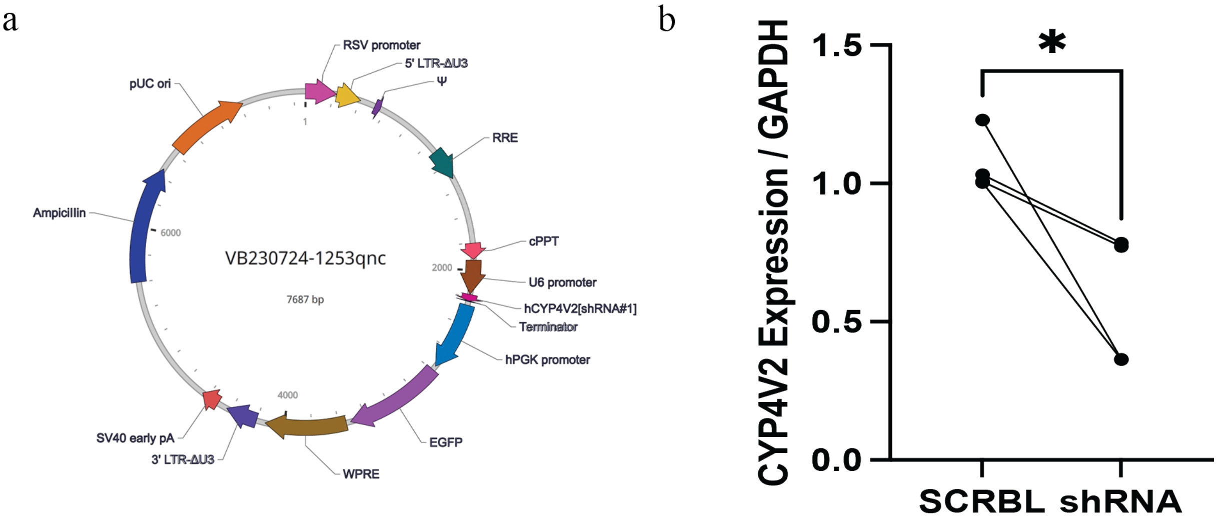


**Suppl. Fig. 8: Validation of CYP4V2 knockdown.** A) CYP4V2 shRNA or SCBL shRNA lentivirus vector. B) Validation of CYP4V2 KD with RT-qPCR on WT and mutant neurons, two independent diffentiations and transfections. *$p < 0.05$, Student's t-test.
